# Human placental-derived stem cell therapy ameliorates experimental necrotizing enterocolitis and supports restoration of the intestinal stem cell niche

**DOI:** 10.1101/2020.10.05.327437

**Authors:** Victoria G. Weis, Anna C. Deal, Gehad Mekkey, Cara Clouse, Michaela Gaffley, Emily Whitaker, Jared A. Weis, Marshall Z. Schwartz, Anthony Atala

## Abstract

Necrotizing enterocolitis (NEC), a life-threatening intestinal disease, is becoming a larger proportionate cause of morbidity and mortality in premature infants. To date, therapeutic options remain elusive. Based on recent cell therapy studies, we investigated the effect of a human placental-derived stem cell (hPSC) therapy on intestinal damage in an experimental NEC rat pup model. NEC was induced in newborn Sprague-Dawley rat pups for 4 days via formula feeding, hypoxia, and LPS. NEC pups received intraperitoneal (ip) injections of either saline or hPSC (NEC-hPSC) at 32 and 56 hours into NEC induction. At 4 days, intestinal macroscopic and histological damage, epithelial cell composition, and inflammatory marker expression of the ileum was assessed. Breastfed (BF) littermates were used as controls. NEC pups developed significant bowel dilation and fragility in the ileum. Further, NEC induced loss of normal villi-crypt morphology, disruption of epithelial proliferation and apoptosis, and loss of Paneth cells and LGR5+ stem cells in the crypt. hPSC treatment improved macroscopic intestinal health with reduced ileal dilation and fragility. Histologically, hPSC administration had a significant reparative effect on the villi-crypt morphology and epithelium. In addition to a trend of decreased inflammatory marker expression, hPSC-NEC pups had increased epithelial proliferation and decreased apoptosis when compared to NEC littermates. Further, the intestinal stem cell niche of Paneth cells and LGR5+ stem cells was increased with hPSC therapy. Together, these data demonstrate hPSC can promote epithelial healing of NEC intestinal damage in part through support of the intestinal stem cell niche.

**New and Noteworthy:** These studies demonstrate a human placental-derived stem cell (hPSC) therapeutic strategy for necrotizing enterocolitis (NEC). In an experimental model of NEC, hPSC administration improved macroscopic intestinal health, ameliorated epithelial morphology, and supported the intestinal stem cell niche. Our data suggest that hPSC are a potential therapeutic approach to attenuate established intestinal NEC damage. Further, we show hPSC are a novel research tool that can now be utilized to elucidate critical neonatal repair mechanisms to overcome NEC disease.

## Introduction

Necrotizing enterocolitis (NEC) is a life-threatening intestinal disease in premature infants. Correlating with the degree of prematurity, NEC affects 5-10% of premature infants and continues to be a leading cause mortality in premature infants (19, 32). Decades of research have shown NEC is a multi-faceted disease that results from the complex interaction of early bacterial colonization, an exaggerated inflammatory response, and immature intestinal epithelium. The premature intestine with an immature immune system can become hyperinflammatory in response to early bacterial colonization (48). In human and animal studies, NEC pathogenesis is accompanied with a significant influx of pathogenic immune cells and increased inflammatory cytokine signaling (15, 39, 47, 72). Early bacterial colonization can also directly damage the epithelial barrier of the intestine. Uniquely upregulated in the premature intestinal epithelium, Toll-like receptor 4 (TLR4), a recognition receptor for gram-negative bacterial endotoxin LPS, is a key initiator of NEC pathogenesis (3). Bacterial LPS activates TLR4 in the premature epithelium and triggers increased enterocyte cell death coupled with decreased proliferation of intestinal stem cells that results in the loss of a stable epithelial barrier (51). Additionally, Paneth cells located in the crypts serve a central role in supporting intestinal stem cell niche homeostasis and in protecting against pathogens (59, 66). As recent studies have shown, loss of Paneth cells can also contribute to NEC pathogenesis (45, 60, 80). Together, these epithelial effects lead to barrier dysfunction and impaired regeneration. The neonatal endogenous repair processes are not able to counter the resultant cyclic damage of bacterial translocation, uncontrolled inflammation, and epithelial damage. In this milieu, the characteristic NEC features can develop including feeding intolerances, bloody stool, and distended and necrotic intestine, eventually leading to systemic sepsis. Although our understanding of NEC instigating factors and potential protective strategies continues to advance, treatment options for established NEC disease have remained relatively unchanged.

As known instigating factors, the microbiome, immune response, and epithelial development have been the focus of preventative and therapeutic investigations in NEC disease. Breastmilk has been shown to protect against NEC-instigating pathways and reduce NEC incidence in experimental NEC models and in premature infants (26, 34, 64). Recently, amniotic fluid has demonstrated similar affects experimentally (25). Accordingly, these fluids contain bioactive factors that inhibit TLR4-mediated epithelial injury, reduce inflammatory signaling integral to NEC pathogenesis, and support intestinal maturation (25, 26). Similarly, probiotic administration has yielded promising results in establishing a healthy microbiome, supporting epithelial development, and reducing NEC incidence (28, 50). These protective pathways have led to new potential prophylactic strategies to reduce the risk of NEC onset. However, they alone often fail to resolve already established NEC damage and therefore are currently limited to prevention of onset (56). Currently, no effective treatments that can heal established NEC disease are available.

Emerging perinatal stem cell approaches from our group and others have been shown to possess dual supporting roles in modulating pathogenic inflammation and resolving epithelial damage through multiple regenerative pathways (49, 55, 69, 77). In experimental NEC, perinatal stem cells can decrease inflammatory signaling, improve epithelial function, and reduce NEC damage (13, 44, 77). As few of the perinatal stem cells were observed to engraft into the intestine in these studies, the beneficial effects were exerted predominantly via paracrine signaling (77). Further, perinatal stem cell conditioned media or exosomes via various administration routes can reduce NEC damage and mortality in experimental NEC (43, 77). Previous perinatal cell studies have focused on stem cells derived from amnion or amniotic fluid obtained mostly via amniocentesis procedures. However, the placenta, long considered medical waste that can simply be collected after healthy birth, is now being regarded as a more optimal tissue for perinatal stem cell isolation. hPSC possess similar proliferative and multipotency characteristics to other perinatal stem cells, however, they also express a unique secretome (41). hPSC have been shown to have immunomodulatory factors and capabilities (40), and also express several factors that can directly promote intestinal epithelial healing and regeneration, including ICAM1, NRG1, EGFR, and Lactoferrin (41). Together, these data suggest that hPSC may be a potential multifaceted therapy to combat NEC pathogenesis.

In this study, we hypothesized that hPSC administration after the onset of NEC induction would ameliorate the established intestinal damage in an experimental model of NEC through immunomodulation and attenuation of epithelial disruption. Expanding upon previous NEC neonate rat models that show NEC damage starting at 24 hours (6, 58, 79), we increased the LPS doses during the first 30 hours of NEC induction to allow the disease to progress further prior to the first hPSC injection. This adjusted timetable allows for more precise investigation of hPSC for therapeutic versus prophylactic use. Here, we evaluated the efficacy of hPSC treatment on survival, clinical symptoms, gross intestinal health, and cellular ileal damage including inflammatory signaling and epithelial cell composition. Using our modified NEC model, our results show hPSC that administration can promote intestinal healing of NEC damage, in part by re-establishment of the intestinal stem cell niche. These findings also have important implications in providing a pivotal research tool that can be leveraged to identify reparative mechanisms that are potential targets in combating NEC disease.

## Materials and Methods

### Placental Cell Isolation, Expansion, and Culture

Consent for obtaining human placenta was obtained from patients prior to giving birth. The Manufacturing Development Center within the Wake Forest Institute for Regenerative Medicine (WFIRM), Wake Forest School of Medicine, collected the full-term placenta specimens. From the whole placental tissue, the chorion was biopsied and then digested to isolate perinatal placental cells as previously described (11, 41). Briefly, isolated placental cells were plated in α-MEM supplemented with AmnioMax and Glutamax. After cell expansion, cell selection for C-kit positive cells was performed with a CD117 antibody (Cat # 120-099-672, Miltenyi Biotec, Germany) in a Miltenyi Mini Macs System. Selected C-kit positive placental cells (hPSC) were expanded in culture and underwent sterility, endotoxin, mycoplasma, and Karyotype testing. Further, phenotypic testing of placental cell cultures confirmed CD29(+), CD44(+), CD73(+), CD105(+), CD146(+), SSEA-4(+), HLA-ABC(+), CD34(-), CD45(-), and HLA-DR(-). Cells were cryopreserved in CryoStor10 until further use. For NEC studies, hPSC between passage 8-12 were cultured to no more than 70% confluency. hPSC were harvested with TrypLE (ThermoFisher, MA), washed twice with DPBS, and diluted to 2×10^6^ cells per 50 µl in DPBS for injection.

### NEC Animal Model

The care, maintenance, and treatment of animals in these studies adhere to the protocols approved by the Institutional Animal Care and Use Committee of Wake Forest University. Timed-pregnant Sprague-Dawley rats were obtained from Charles River Laboratories and monitored for birth beginning 36 hours prior to expected birth. Within 6 hours of birth, newborn pups from each litter were randomly divided into 3 groups: breastfed (BF) controls (n=11), NEC (n=11), and NEC with subsequent hPSC therapy (NEC-hPSC, n=11). BF controls were left with dam and received no additional stressors. NEC pups were separated from dam and housed in a Caleo Infant Incubator (Drager, PA) at 30°C with 50% humidity for the remainder of the study. NEC was induced with modifications of an established protocol (5, 56, 77, 79). NEC pups were oral gavage feed via a 22ga plastic feeding tube (Instech Laboratories, PA) 4 times a day with a hyperosmolar formula prepared from 15 g Similac PM 60/40 (Abbott Nutrition, OH) in 75 mL Esbilac canine supplement (Pet-Ag, IL). NEC pups also received oral administration of LPS (Cat #L3012, Sigma, MO; 4 µg/g, 3X within 30 hours of birth), and hypoxia (5% O_2_ for 10 minutes, 3X/day) (Figure 1). NEC pups each received 2 doses of either 50 µl DPBS (NEC group) or 2×10^6^ hPSC in DPBS via intraperitoneal (IP) injection. Timed for after initial NEC damage onset(6, 58, 79), the first hPSC injection was given after the last of the 3 LPS doses (32 hours of NEC induction) with a second hPSC injection 24 hours later (56 hours). A subset of pups also received an IP injection of EdU (50 mg/kg, Cayman Chemical, MI) 24 hours prior to euthanasia. To avoid previously observed fluid leakage from the injection site, all injections were performed under light anesthesia with pups receiving similar doses and lengths of anesthesia (<10 minutes of Isoflurane). At 96 hours of NEC induction, all remaining pups were euthanized.

**Figure 1.**
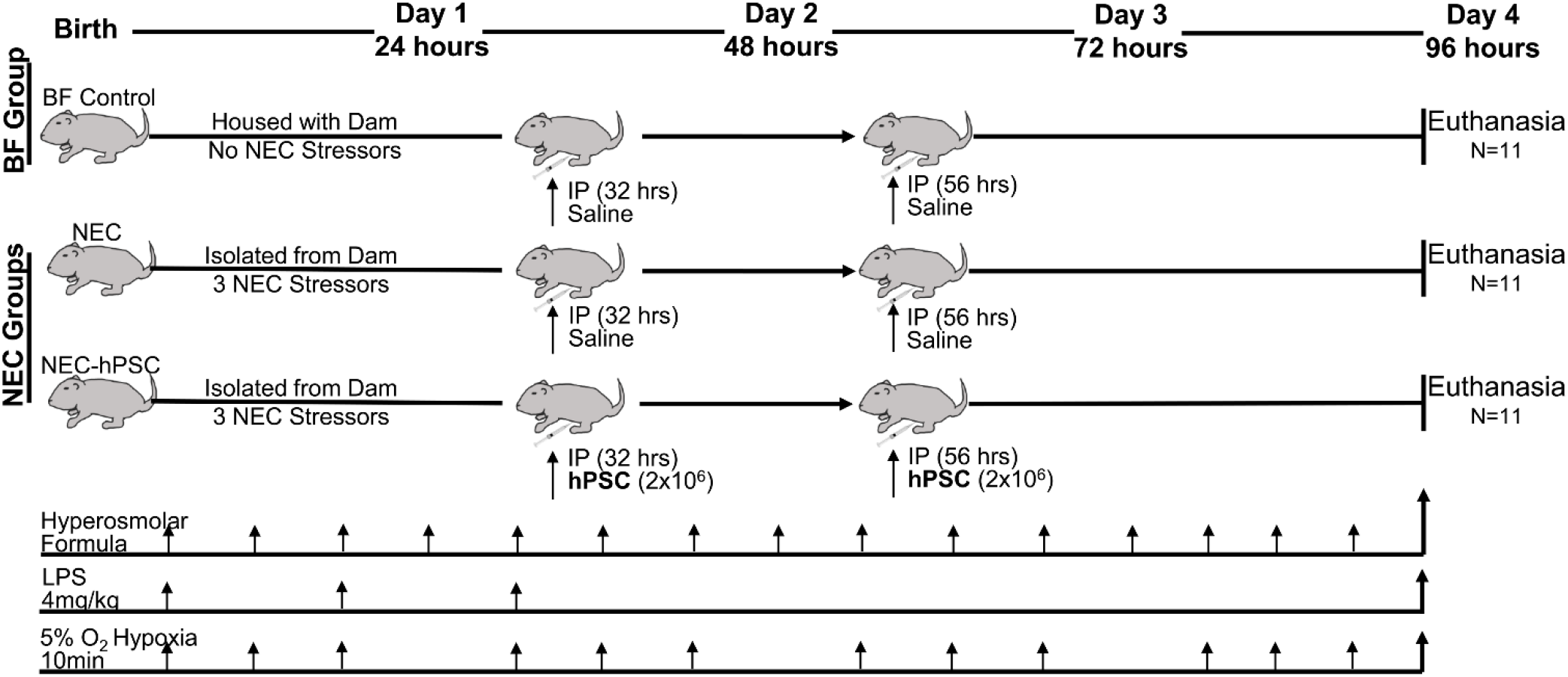
Study design schematic. Newborn Sprague-Dawley rats were randomly divided into 3 groups: BF control, NEC, NEC-hPSC. Within 6 hours of birth, NEC and NEC-hPSC pups began receiving the 3 NEC stressors of hyperosmolar formula feeding, LPS oral dosing, and whole animal hypoxic stress. After administration of the 3 LPS doses, NEC pups received an initial IP injection of saline (NEC) or hPSC (NEC-hPSC) at 32 hours with a second IP injection 24 hours later. Body weight and Clinical Sickness Score was assessed daily. All remaining pups were euthanized at 96 hours resulting in N=11 for all experimental groups.

### Survival, Weight, and Clinical Sickness Score

Pup death or humane euthanasia up to the 96-hour endpoint was assessed. Pup deaths caused by known or suspected adverse technical events (perforation, formula aspiration) were removed from the survival analysis (n=2). Body weight and a Clinical Sickness Score were assessed at birth and every 24 hours. Due to variations in birth weights between litters, body weight was analyzed as the daily change in weight to minimize litter to litter variations. The Clinical Sickness Score, as previously described (77, 79), provides a composite score from 4 selected criteria: Appearance, Response to touch, Natural activity, and Body color. A two-way ANOVA was performed to identify interactions between the individual variables. Statistical significance (*p*<0.05) between groups at each time point was determined using the Mann-Whitney *U* two-tail test.

### Macroscopic Score and NEC Grade

Following euthanasia, the entire intestine (duodenum-colon) was excised with removal of its surrounding and connective tissues, straightened, and photographed. A Macroscopic Score was assessed based on consistency/fragility, color, and degree of dilatation as previously described (77, 79). Extent of dilation and severity was further assessed by a custom-built software that allows measurement of diameter along the length of the intestine. Images were processed by semi-automatic segmentation and image morphological operations to yield a region-of-interest (ROI) mask of the intestine using an active contour-based region growing algorithm (42) to automatically refine a manually designated initial ROI. The final segmentation of each intestine was then manually reviewed and edited, if necessary, to ensure optimal segmentation. Next, the diameter along the length of the intestine was calculated using a Euclidean distance transform of the masked ROI. Distances were calibrated using a reference scale that was photographed within the same image field as the intestine. The medial ileal diameter for each animal was then calculated using diameter measurements in 100 µm increments along the length of the ileum with the ileum designated as the final third of the total intestinal length. To determine the extent of pathological involvement of the ileum, the median ileal diameter in the BF control group was used as the healthy dilation control. Pathological distention was then defined as an ileal diameter (within the 100 µm increment) greater than the BF median diameter plus 2 standard deviations. The percentage of pathological involvement for each animal was calculated by determining the length of the ileum defined as pathologically distended divided by the total length of the ileum. Length of small intestine was also assessed. Statistical significance (*p*<0.05) was performed between the groups using the Mann-Whitney *U* two-tail test.

For histological assessment, the small intestine was fixed in 4% paraformaldehyde overnight at 4°C and embedded in paraffin. Sections were deparaffinized, rehydrated, and hematoxylin and eosin (H&E) staining was performed. Based on a previously established scoring system to assess NEC damage (77, 79), H&E stained ileum was evaluated for an overall NEC Grade: 0 (normal), 1 (disarrangement of villus cells, mild villus core separation), 2 (disarrangement of villus cells, severe villus core separation), 3 (epithelial sloughing), 4 (bowel necrosis/perforation). NEC was defined as grade 2 or above. Ileal H&E sections were then imaged at 10X with an EVOS FL Auto 2 Cell Imaging System (Invitrogen, CA) to measure extent of ileal involvement of each NEC Grade. For this, the linear length of ileum with each NEC Grade was measured and then normalized by the total linear length of the ileum assessed (% involved = length_Grade_ / total ileal length). Statistical significance (*p*<0.05) between the groups was determined using the Mann-Whitney *U* two-tail test.

### qPCR Analysis

To examine inflammatory signaling, a small section of distal ileum was frozen for RNA isolation. RNA was extracted with TRIzol (Invitrogen) according to the manufacturer’s instructions. The RNA (1 µg) was treated with RQ1 RNase-free DNase (Promega, WI) and then reverse-transcribed using High-Capacity cDNA RT kit (Applied Biosystems, CA). Equal amounts of each cDNA were analyzed by real-time PCR with specific primers and PowerUp SYBR Green (Invitrogen) in an ABI QuantiStudio 3 real-time PCR system. Each sample was measured in triplicate. The following pre-designed primers from IDTDNA (CA, USA) were used: TNFα forward: 5’- GTC TTT GAG ATC CAT GCC ATT G-3’, reverse: 5’- AGA CCC TCA CAC TCA GAT CA-3’; IL1β forward: 5’- TTG TCG TTG CTT GTC TCT CC-3’, reverse: 5’- GTG CTG TCT GAC CCA TGT-3’; NFκB forward: 5’- GAC TCT TCT TCA TGA TGC TCT TG-3’, reverse: 5’- GAG TTC CAG TAC TTG CCA GAC-3’; and TBP forward: 5’- GGA GAA CAA TTC TGG GTT TGA TC-3’, reverse: 5’- TGT GAA GTT CCC CAT AAG GC-3’. As previously described (71), cycle threshold was converted to relative expression via the 2^-ΔΔ^ cycle threshold method, using TATA-box-binding protein (TBP) as an endogenous control. For each analysis, the mean value of the normalized cycle thresholds of all breastfeed rat samples was used as reference. With less than 10 samples for some groups due to tissue degradation/poor RNA quality, statistical analysis of qPCR was performed as a one-tail test with the hypothesis that NEC had increased inflammatory expression and NEC-hPSC was decreased compared to untreated NEC. Statistical significance (*p*<0.05) between the groups was determined using the Mann-Whitney *U* test.

### Immunostaining and Quantifications

Paraffin sections were deparaffinized and rehydrated before antigen retrieval was performed using Antigen Retrieval Buffer (Abcam, MA) in a pressure cooker for 15 minutes. After cool-down on ice, sections were blocked with protein block serum-free (Dako, Denmark) for 1 hour at room temperature. The following primary antibodies were incubated at 4°C overnight: Rabbit anti-Cleaved Caspase 3 (Cat# 9661, Cell Signaling Technologies, MA), Rabbit anti-OLFM4 (Cat# 39141T, Cell Signaling Technologies), Mouse IgG_2b_ anti-Ezrin (Cat# CPTC-Ezrin-1, Developmental Studies Hybridoma Bank, IA), and Rabbit anti-Lysozyme (Cat# MBS2556232, MyBioSource, CA). After 3 washes in 1X phosphate-buffered saline for 5 minutes each, sections were incubated for 1 hour at room temperature with the appropriate secondary antibodies conjugated for immunofluorescence with Alexa Fluor 488, Cy3, or Cy5 (1:1000; Jackson ImmunoResearch, PA). Slides were then washed and incubated with 4’, 6-diamidino-2-phenylindole (DAPI, 1:10,000) for 5 minutes. After additional washing, slides were mounted with ProLong Gold Antifade Reagent (Invitrogen). For EdU dectection, Click-iT EdU Proliferation Kit for Imaging was used (Invitrogen). Briefly, after antigen retrieval, slides were blocked with 3% BSA, permeabilized with 0.5% Triton X-100, and EdU labeled followed the manufacturer protocol. Co-immunostaining then proceeded to protein block as described above. Sections were imaged using an Olympus FV3000 fluorescent microscope (Olympus) equipped with a Nuance EX camera (PerkinElmer) or EVOS FL Auto 2 Cell Imaging System (Invitrogen).

Ezrin immunostaining (with phase contrast as needed) was used for villus/crypt height quantification. From each animal, 3 images were taken of randomly selected regions of well-oriented ileum. Well-oriented crypts and villi were identified in each 20X field image and the height of each was measured using FIJI. The villi were measured from the villus tip to the crypt transition and the crypts were measured from the crypt transition to the bottom of the invagination between two villi. From Cleaved Caspase 3 (apoptosis), EdU (proliferation), OLFM4 (intestinal stem cell), and Ezrin (enterocyte) immunostainings, 3 randomly selected 20X field images from each animal were quantified as previously described (70). Briefly, positively stained cells within the epithelial layer, the top of villi, and the base of crypts were manually marked in individual “layers” using Photoshop (Adobe Systems, CA). Analysis of total positive-cell numbers and quantitative spatial localization along the crypt-villi axis was conducted with a modified custom-built MATLAB software (Mathworks, MA) previously described (70). Total cell counts were normalized to the linear length of ileal tissue quantified as automatically measured from the ‘crypt base’ contour line. Histograms with bin widths equal to 25 pixels (10 µm) were generated for each marker expression. Cell counts in histogram bins were normalize to the total number of crypts manually counted in each animal to reflect the average number of cells per crypt that are localized within each bin location. Histograms are reported as a mean histogram line with standard deviation designated as a shaded region. Similar total cell counts and distributions were found between normalization to crypt number and linear length. As Paneth cells are not as abundant in 4 day-old rat pups, larger regions of Lysozyme-immunostained ileal sections (4-8 mm for each animal) were scanned with an EVOS system (Invitrogen) and the total number of Lysozyme-positive epithelial cells per linear ileal length was manually quantified using FIJI. Although initial investigation showed minimal differences in the number of crypts per mm of ileum, normalization to ileal length for total cell counts was chosen to minimize any potential difficulties in crypt identification in NEC-damaged tissue and therefore allow for more high-throughput analysis. For the N<10 sample size used in EdU subset analysis, Mann-Whitney *U* one-tail test was used with the hypothesis that NEC had reduced cell numbers than BF and NEC-hPSC had greater cell numbers than NEC. For all other cell counts, statistical significance (*p*<0.05) between the groups was determined using the Mann-Whitney *U* test.

## Results

### The NEC animal model was successfully replicated with increasing insult prior to treatment and decreased survival in all experimental groups

Beginning within 6 hours of birth, neonatal rat NEC models utilize formula feeding, LPS administration, and hypoxic stress to induce NEC (79). Previous investigations have established that intestinal damage is initiated within 24 hours of NEC induction, while overall mortality limits the study window to 96 hours (6, 24, 58, 79). To increase intestinal insult prior to our treatment timepoint, the NEC rat pup model used in the present study included 3 doses of LPS within the first 30 hours of NEC induction (Figure 1) in comparison to 1-2 LPS doses of previous experimental NEC studies (44, 77, 79). Accordingly, pups undergoing NEC stressors showed body weight loss and increased Clinical Sickness Scores within 24 hours and these symptoms continued to worsen until the 96-hour endpoint (Figure 2A, B). Breast fed (BF) controls gained weight from birth to endpoint and showed no significant clinical symptoms for the duration of the study.

**Figure 2.**
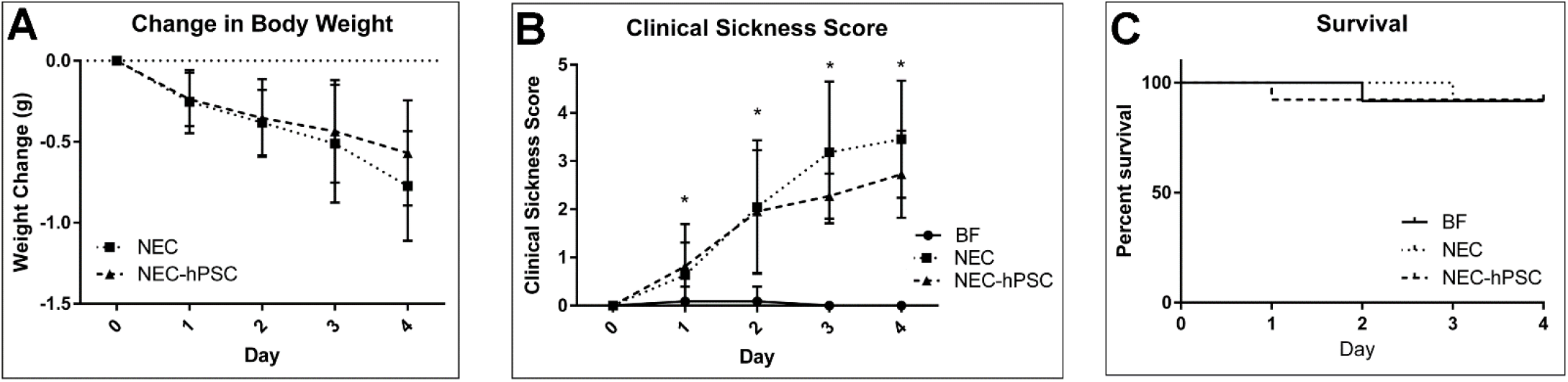
Whole animal assessments of mortality and symptoms. (A) NEC and NEC-hPSC animals had noticeable weight loss beginning within 24 hours prior to initial IP injections. Both groups continued to lose weight until study conclusion. While NEC-hPSC animals appeared to trend towards slower weight loss in the last 48 hours, a statistical difference was not reached between weight loss changes of the 2 groups. (B) A composite score called Clinical Sickness Score was used to assess clinical symptoms and overall health of the animal (0=healthy, normal). BF animals show no clinical symptoms. NEC and NEC-hPSC animals have increased symptoms by Day 1 (24 hours) compared to BF controls (\**p*=0.02). The Clinical Sickness Scores of NEC and NEC-hPSC animals are identical until Day 2 when NEC-hPSC scores begin trending lower. However, no statistical difference is observed at the study endpoint of 96 hours. (C) No significant difference was observed in survival. Overall, each group lost 1 animal over the course of the 4-day study. The animals lost prior to the 96-hour endpoint were not included in further analyses.

To study hPSC therapeutic potential, two injections of 2×10^6^ hPSC were given 24 hours apart following completion of LPS administration (Figure 1). These injection timepoints strategically occurred after the previously reported intestinal damage initiation at 24 hours and the observed onset of clinical symptoms in the present study (Figure 2A, B). Prior to hPSC administration, NEC-hPSC pups and untreated NEC pups showed similar body weight loss and increased Clinical Sickness Scores. After the 2^nd^ hPSC injection, body weight loss and Clinical Sickness Scores began trending towards improvement in NEC-hPSC pups, however these trends did not reach statistical significance by the 96-hour endpoint (Figure 2A, B). No difference between the groups was observed in survival at Day 4 (Figure 2C).

### hPSC treatment improved gross intestinal pathology

Distended and hemorrhagic intestine leading to complete necrosis are characteristic symptoms of NEC pathogenesis. As these pathologies are predominantly found in the ileum and not commonly found in the duodenum in NEC patients or experimental NEC models, we focused our analysis on the ileum in our studies. Upon gross ileal examination in NEC pups, obvious pathologies of dilation and tissue fragility were observed (Figure 3 and Supplemental Figure 1). A previously established composite score for dilation, coloration (from hemorrhage or necrosis), and consistency (tissue fragility) (79) was used to assess these gross pathological symptoms on a scale of 0 to 2 (Figure 3B). The NEC induction procedures increased the ‘Macroscopic Score’ of the ileum from NEC pups compared to BF controls (NEC: 1.18; BF: 0.06). However, NEC pups treated with hPSC had a significantly reduced gross pathology composite score compared to untreated NEC ileum (NEC-hPSC: 0.76). Examination of the individual categories of the ‘Macroscopic Score’ revealed that all 3 components contributed in part to the increased score in NEC pups. Consistency and dilation, but not coloration were the score-driving factors for the reduced disease pathology in NEC-hSPC pups (Supplemental Figure 1A, B, C).

**Figure 3.**
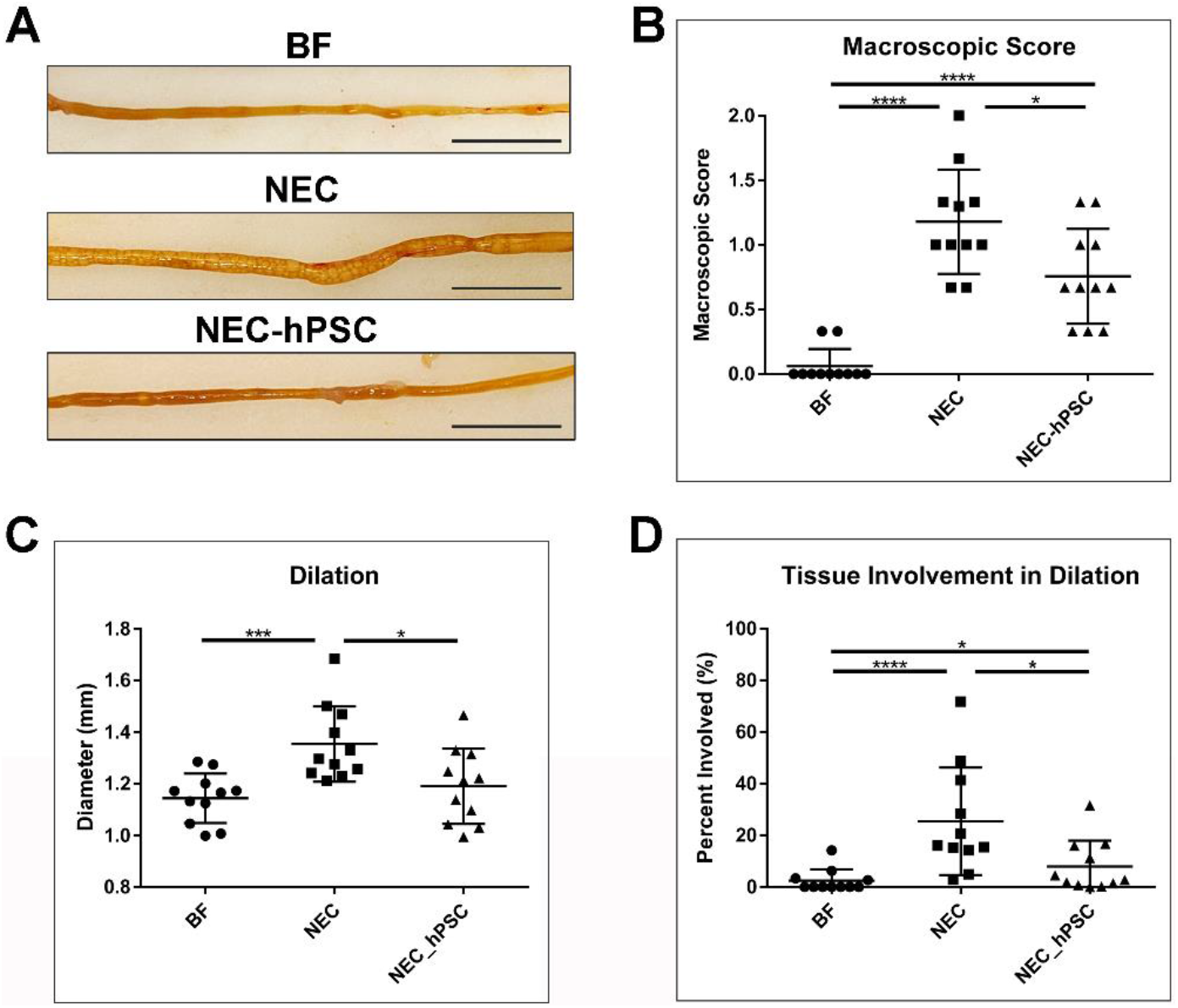
Macroscopic health of ileum. (A) In representative images of ileum, NEC animals have significant distention that can affect bowel function compared to BF control animals. This distention is reduced in NEC-hPSC animals. (scale bar = 1 cm) (B) An established composite Macroscopic Score that included coloration, consistency, and dilation was used to assess the overall gross pathological health of the ileum (0=healthy, normal). NEC animals have a significantly worse Macroscopic Score compared to BF controls (NEC=1.18±0.40; BF=0.06±0.13, \*\*\*\**p*<0.0001). hPSC treatment reduced the macroscopic pathology of the ileum in NEC-hPSC animals (NEC-hPSC=0.76±0.37, \**p*=0.031). (C) As the key driving factor of the Macroscopic Score, the severity of ileal dilation was quantified by measuring diameter along the entire length of the ileum. The median of the ileal diameters shows a significant 11% increase (0.13 mm) in NEC animals (NEC=1.30±0.146 mm; BF=1.17±0.096 mm, \*\*\**p*=0.0003). Ileal dilation in NEC-hPSC animals is decreased back to BF levels (NEC-hPSC=1.21±0.146 mm, \**p*=0.0193). (D) To assess the amount of tissue involved, the percent of the ileal length with significant pathological distention was measured. NEC animals had almost a 10.5-fold increase of tissue with pathological distention compared to BF control animals (NEC=25.4%±20.8; BF=2.42%±4.43, \*\*\*\**p*<0.0001). hPSC treatment of NEC animals decreased tissue involvement to less than 10% of the ileum (NEC-hPSC=7.95%±10.0; \**p*=0.011 vs NEC and \**p*=0.036 vs BF).

As excessively distended intestine can cause impaired motility and predispose the intestine to dysbiosis and malabsorption (10, 12, 33), we sought to further investigate the severity and extent of the ileal dilation in experimental groups. Furthermore, the general stunting of tissue growth that can occur in artificially-reared rat pups (63, 75) could potentially bias one’s perception of ileal dilation severity and result in a skewed ileal dilation scoring for the composite ‘Macroscopic Score’. Therefore, a custom software was developed to automate measurements of the exact ileal diameter along the length of the tissue. This method allowed for a quantitative analysis of the severity of the characteristic NEC intestinal distention and provided the ability to examine the amount of ileal length that presented this distended NEC pathology. In NEC pups, the ileum was significantly dilated compared to BF controls (NEC: 1.3 mm, BF: 1.17 mm) (Figure 3C). hPSC treatment in NEC pups alleviated ileal distention back to BF levels (NEC-hPSC: 1.21 mm). These findings help establish an objective quantification method to measure the severity of NEC-associated ileal distention, a leading factor in previously used gross pathological composite scoring. Additionally, as NEC pathologies can range from localized patches to total ileal involvement, this novel quantification method was also used to assess the amount of ileal tissue that had this pathological dilation (Figure 3D). Pathological dilation of the ileum was defined as a diameter greater than the BF median diameter plus 2 standard deviations (95% confidence interval). In our experimental NEC model, greater than 25% of the ileum was involved with this gross distention pathology. Upon hPSC treatment, ileal involvement was significantly reduced to less than 10%. Together, these data quantitatively show that hPSC can reduce both the severity of ileal distention and the extent of tissue involvement.

### Histological NEC damage is ameliorated with hPSC administration

H&E stained sections of the ileum were analyzed for histopathological scoring of damage on a scale of 0 (normal) to 4 (transmural necrosis) (79). An overall histological score of 2 and above was considered NEC. No morphological changes were observed in BF control pups (NEC Grade: 0) (Figure 4). In NEC pups, the epithelial layer was disrupted with loss of crypt morphology and varying degrees of cellular and villar sloughing, indicative of NEC damage (Figure 4A). Occasional, localized segments displayed red blood cells within the lumen. After 4 days of NEC stressors, 10 out of 11 neonates developed NEC for a mean NEC Grade of 2.0 (Figure 4B). hPSC treatment significantly reduced the NEC-induced damage. Upon hPSC administration, a re-emergence of normal crypt/villi morphology and decreased epithelial disruption was observed (Figure 4A). NEC-hPSC animals had an improved NEC Grade of 1.41 and reduced overall NEC incidence of 9% (Figure 4B).

**Figure 4.**
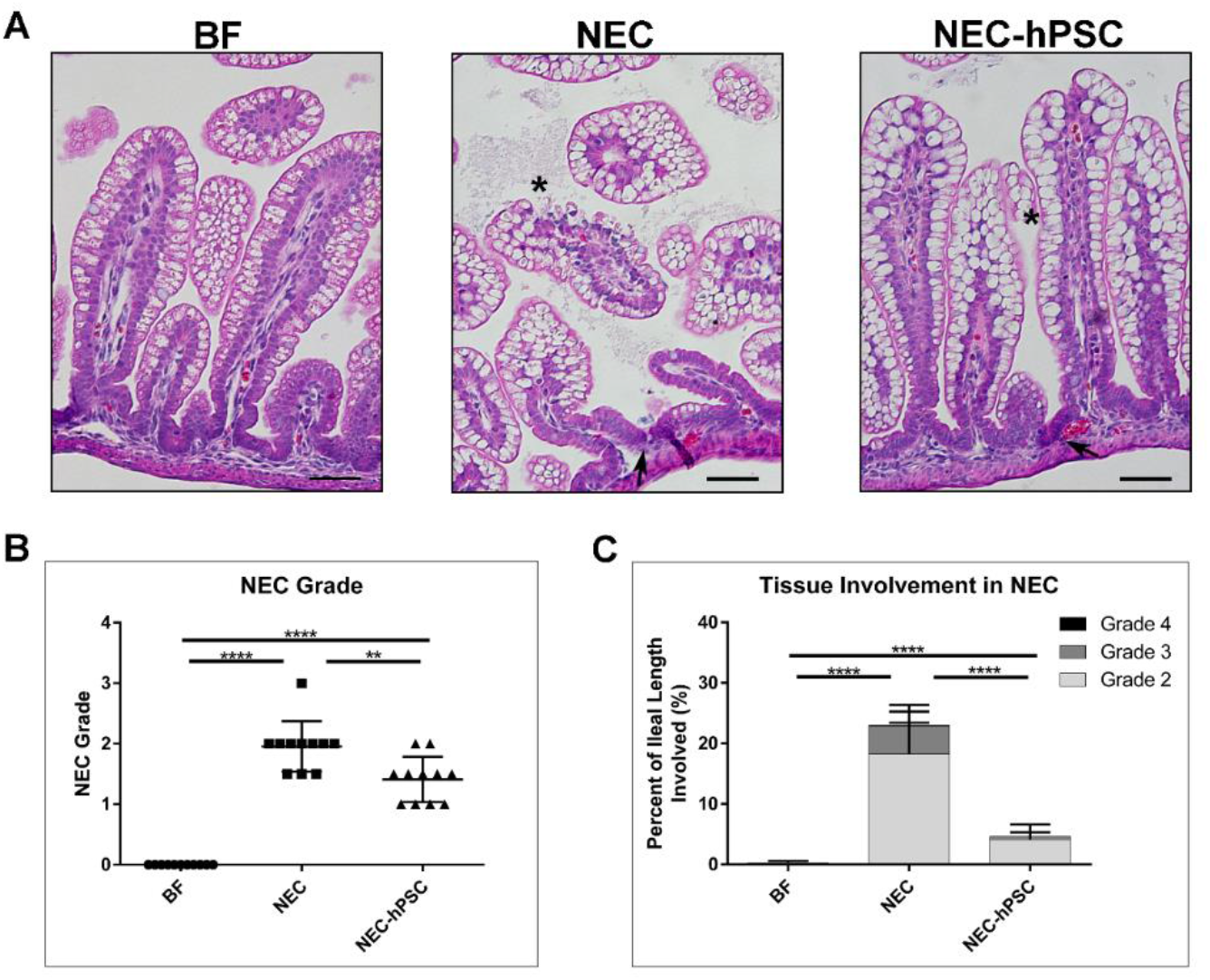
Histological damage in NEC. (A) H&E stained ileal sections show normal histology in BF control animals. NEC animals have loss of normal crypt architecture (arrow), disrupted epithelial cell morphology with cellular sloughing (asterisk), and some villar sloughing. Crypt morphology begins to re-emerge in NEC-hPSC animals (arrow). Villi appear more intact with resolving epithelial cell order (asterisk). (scale bar = 50 µm) (B) NEC associated damage in each ileum was assessed using an established histological scoring and given an overall NEC Grade. An NEC Grade ≥2 was considered NEC. NEC animals had an increased NEC Grade of 2.0±0.416 compared to BF animals’ score of 0.0±0.0 (\*\*\*\**p*<0.0001). hPSC treatment of NEC significantly decreased histological NEC damage to 1.41±0.38 (\*\**p*=0.0074 vs NEC). (C) The amount of ileal tissue with each of the NEC Grades of damage was assessed. Over 20% of the ileum from NEC animals displayed NEC Grade ≥2 with almost 5% being the more severe NEC Grade 3. Compared to NEC animals, NEC-hPSC animals have significantly less tissue with histological damage of NEC Grade ≥2. Less than 5% of NEC-hPSC tissue had NEC Grade 2 damage and only rare occasions of NEC Grade 3 (0.5%). (\*\*\*\**p*<0.0001).

In addition to the severity of histological NEC damage, the amount of intestinal tissue involved in the NEC pathogenesis is also a concern. Therefore, the potential hPSC efficacy in reducing the overall tissue burden was assessed. For each animal, the percentage of ileal length with each Grade of NEC damage (≥2) was measured (% involved = length_Grade_ / total ileal length). On average, approximately 23% of the ileum from each NEC neonate had histological damage of NEC Grade ≥2. The NEC damage was predominately Grade 2 (18.2%) with smaller involvement of Grade 3 (4.6%) and 4 (0.1%) (Figure 4). NEC-hPSC neonates had remarkably reduced ileal involvement with less than 5% of the ileum having histological NEC damage ≥2 with the damage predominately NEC Grade 2 (Figure 4). Furthermore, Grade 3 was rarely observed in NEC-hPSC animals with less than 0.5% ileal involvement and no observed Grade 4. In addition, these histological findings correlate to the extent of ileal distention and the ‘Macroscopic Score’ as previously reported (Supplemental Figure 1) (79). Overall, these histological analyses show significantly reduced severity in NEC damage and tissue involvement in hPSC treated NEC animals compared to untreated NEC.

### hPSC treatment promotes healing through attenuation of epithelial damage

Since hPSC have known immunomodulatory abilities, we sought to investigate the effect of hPSC treatment on inflammation, a hallmark of NEC pathogenesis. Three inflammatory markers previously shown to be associated with NEC were selected to investigate the overall inflammatory milieu in the ileum. In comparison to healthy BF controls, NEC animals had significantly increased expression of TNFα, IL1β, and NFκB (Figure 5A). In NEC-hPSC animals, IL1β, and NFκB showed a decreased trend in NEC-hPSC but did not reach statistical significance (Figure 5A). TNFα was the sole cytokine to be significantly reduced towards the low BF level (Figure 5A). This underwhelming reduction of the prominent inflammatory signaling in NEC-hPSC suggests immunomodulation is not the sole repair pathway employed by hPSC in this ileal setting.

**Figure 5.**
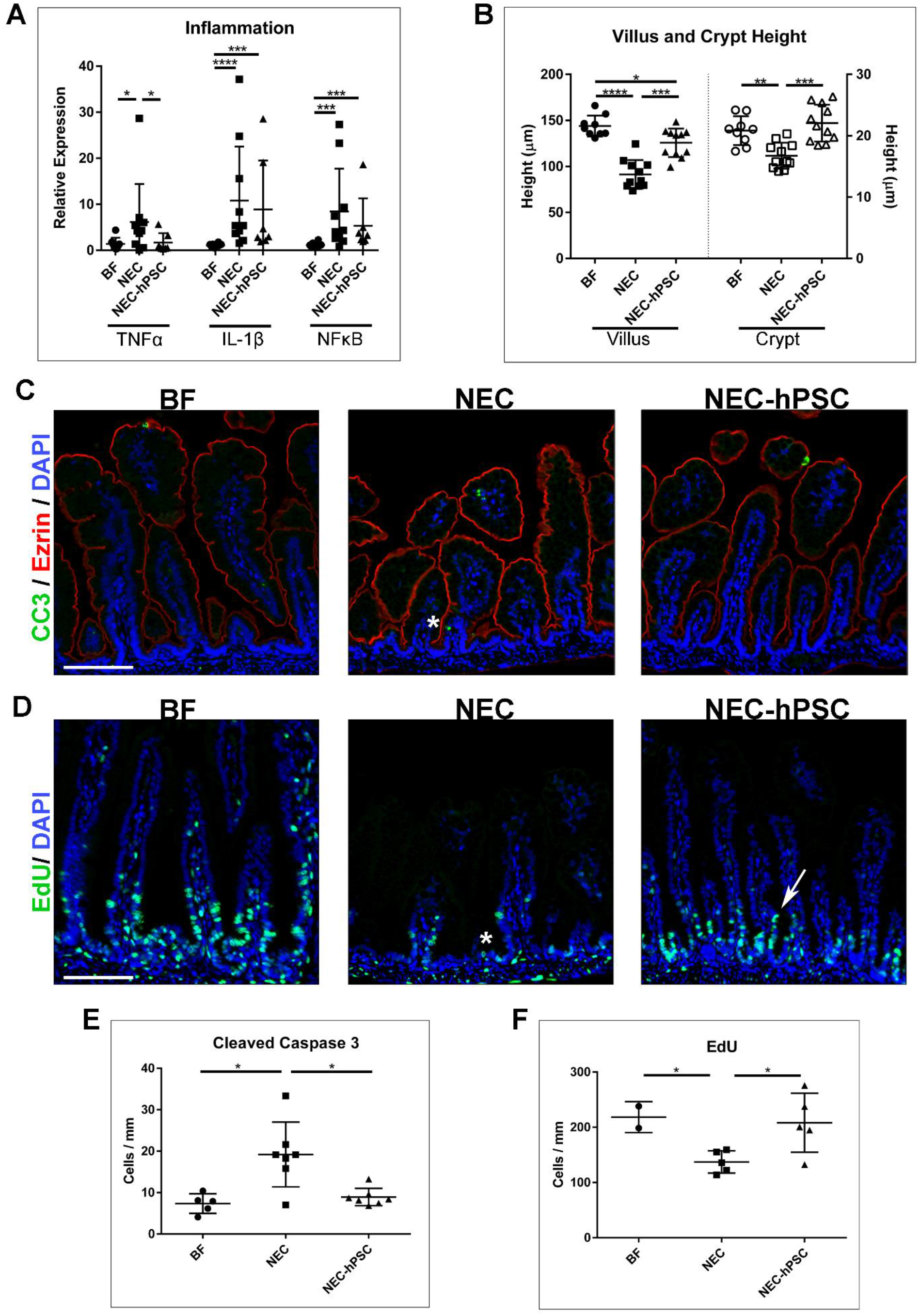
Assessment of inflammation and epithelial barrier. (A) Quantitative PCR for 3 prominent inflammatory signaling biomarkers (TNFα, IL-1β, and NFκB) was performed to examine global inflammatory status. All 3 inflammatory biomarkers were significantly increased in NEC animals compared to BF control animals (NEC: TNFα=6.15±8.25, \**p*=0.034; IL-1β=10.18±11.71, \*\*\*\**p*<0.0001; NFκB=8.44±9.27, \*\*\**p*=0.0007). In NEC-hPSC animals, only TNFα was significantly reduced compared to NEC animals (NEC-hPSC: TNFα=1.69±2.02, \**p*=0.035; IL-1β=8.89±10.61; NFκB=5.31±5.98). IL-1β and NFκB expression was not decreased with hPSC treatment and remained elevated compared to BF animals (\*\*\**p*=0.0002, \*\*\**p*=0.0006, respectively). (B) Villus and Crypt heights were measured to assess overall epithelial status. In NEC, villi and crypts were significant shorter compared to BF controls (NEC: Villus=91.4±15.5 µm, Crypt=16.8±2.1 µm; BF: Villus=143.8±11.5 µm, Crypt=20.8±2.3 µm; \*\*\*\**p*<0.0001, \*\**p*=0.0016, respectively). While NEC-hPSC villar height was not completely restored to BF levels by Day 4 (\**p*=0.031), it was significantly increased compared to NEC animals (125.7±15.4 µm, \*\*\**p*=0.0002). The crypt height in NEC-hPSC was also increased (22.1±3.0 µm \*\*\**p*=0.0001 vs NEC). Notably, NEC-hPSC height was restored back to BF levels. (C) Immunostaining for cleaved caspase 3 (green) showed increased apoptosis in NEC animals (asterisk). This aberrant cell death was not observed in NEC-hPSC animals. (Ezrin=Red, DAPI=blue, scale bar=100 µm) (D) EdU (green) was used to identify proliferation in a subset of animals. NEC animals had marked decrease in proliferating cells emerging from the crypts (asterisk). In NEC-hPSC, normal proliferation and migration began to re-emerge (arrow). (scale bar=100 µm) (E) Quantification of epithelial cells positive for nuclear cleaved caspase 3 shows a significant increase in epithelial cell apoptosis in NEC (19.2±7.8 cells/mm, \**p*=0.0177 vs BF). Aberrant apoptosis is decreased in NEC-hPSC (8.9±2.1 cells/mm, \**p*=0.0175 vs NEC) and reduced to normal BF levels (7.3±2.4 cells/mm). (F) Proliferation denoted by EdU incorporation was quantified in epithelial cells. Proliferation is reduced in NEC (137.2±20 cells/mm, \**p*=0.048 vs BF). Upon hPSC treatment of NEC, epithelial proliferation is significantly increased compared to untreated NEC animals (NEC-hPSC: 208.2±53.44 cells/mm, \**p*=0.028 vs NEC).

The hPSC secretome possesses factors that can potentially support epithelial restitution and repair (41, 55). Therefore, we also examined the epithelial damage in NEC by first assessing the crypt/villus architecture in the ileum. The normal villus and crypt structures found in BF ileum were lost in NEC ileum (Figure 4 and 5B). NEC damage caused shortening of villi and shallow crypt invaginations compared to healthy BF ileum (NEC: Villus= 91.4 µm, Crypt=16.8 µm; BF: Villus=143.8 µm, Crypt=20.8 µm) (Figure 5B). hPSC treatment caused significant lengthening of villi (125.7 µm) (Figure 5B). Moreover, in NEC-hPSC animals, normal crypt depth was increased back to healthy BF control depths (22.1 µm) (Figure 5B). From these combined data, we sought to further investigate the potential reparative effect of hPSC on the intestinal epithelium.

First, we examined epithelial apoptosis in BF, NEC, and NEC-hPSC ileums. Immunostaining for cleaved caspase 3 showed the low homeostatic levels of apoptosis in the epithelium of BF neonates (Figure 5C, E). As has been previously reported (35, 77), the number of apoptotic epithelial cells more than doubled in NEC neonates (Figure 5C, E). In NEC-hPSC animals, epithelial apoptosis was reduced back to BF amounts (Figure 5C, E). We then hypothesized that an inverse trend would be seen for proliferation of the epithelium. To quantify proliferation and cell migration, a subset of animals was intraperitoneally injected with EdU 24 hours prior to euthanasia at the 96-hour endpoint. As previously established in both human and animal model studies, epithelial cell proliferation was decreased in NEC neonates (Figure 5D, F). Compared to the untreated NEC animals, hPSC treated NEC neonates had a significant increase in epithelial proliferation (Figure 5D, F). Together, these data show hPSC reduce cell death and support proliferation within the ileal epithelium.

### hPSC treatment supports enterocytes and the intestinal stem cell niche in NEC repair

Dysfunction and loss of key epithelial cell lineages, specifically enterocytes, Paneth cells, and LGR5^+^ intestinal stem cells, are known to be critically involved in NEC pathogenesis (7, 38, 45). Moreover, our data suggest epithelial support is an important mechanism for hPSC therapy in NEC disease. Therefore, we assessed enterocytes, Paneth cells, and LGR5^+^ intestinal stem cells in each of the experimental animal groups using immunostaining with lineage specific antibodies (Ezrin, Lysozyme, and OLFM4 (67), respectively). Ezrin immunoreactivity was used to quantify enterocytes in the ileum normalized by the linear intestinal length measured (Figure 6A, B). BF controls had normal, ordered enterocytes lining villi with approximately 1013 enterocytes per mm of ileum. In NEC, a decrease of Ezrin-positive enterocytes was observed (701 enterocytes per mm). This loss of enterocytes may be a significant factor in the shorter heights in villi in NEC neonates. As suggested by improved villar height, hPSC treatment significantly increased the number of enterocytes (882 enterocytes per mm), however it did not completely replenish cell numbers back to BF levels.

**Figure 6.**
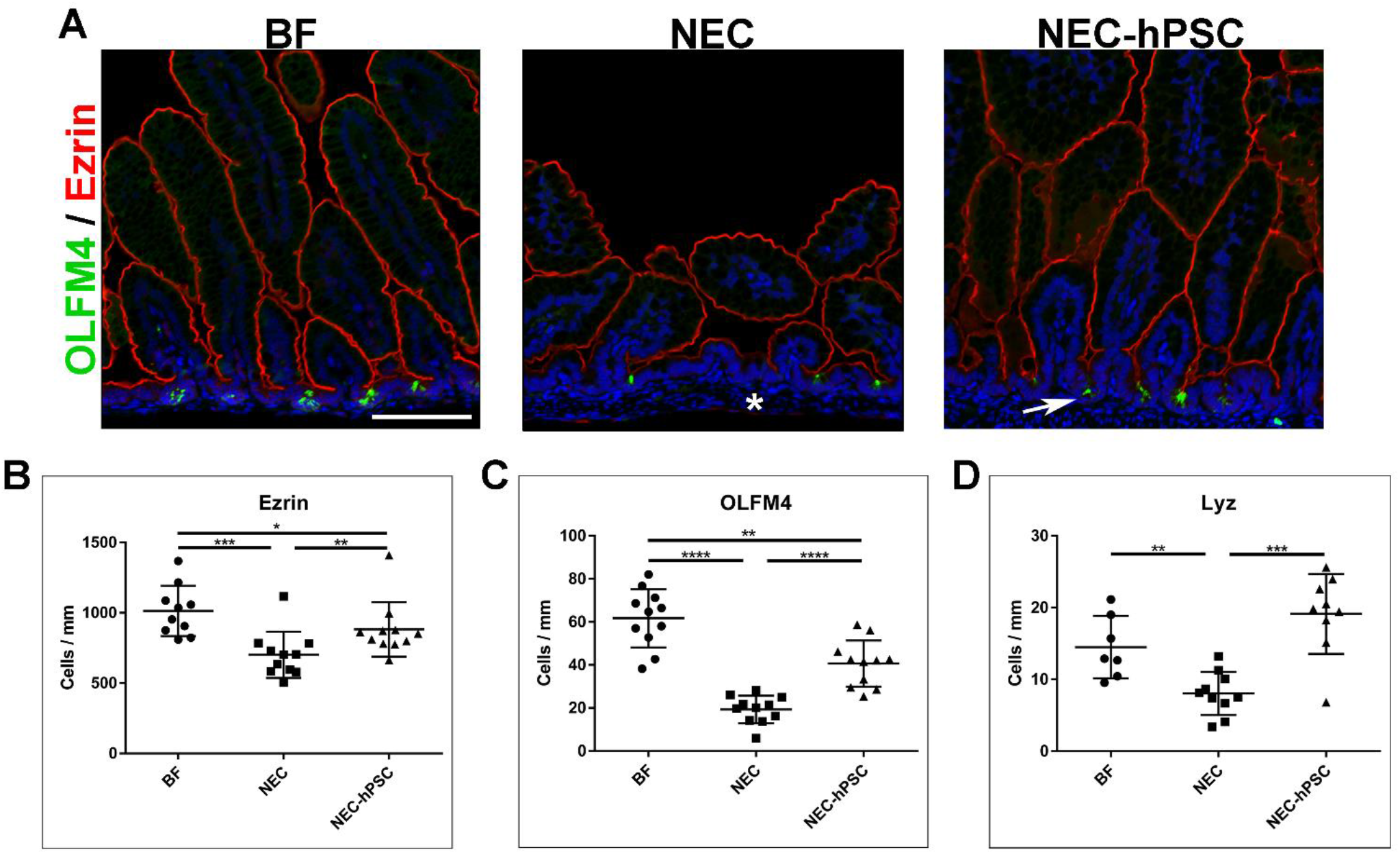
Assessment of epithelial cell lineages. (A) Immunostaining for Ezrin (red) and the LGR5+ intestinal stem cell (ISC) marker OLFM4 (green) illustrated the shortened villi and crypt heights in NEC. Further, it showed a loss of both enterocytes and LGR5+ stem cells (asterisk) in NEC that was partly ameliorated with hPSC treatment (arrow). (DAPI=blue, scale bar=100 µm) (B) Quantification of Ezrin-positive enterocytes showed significant loss in NEC (701.6±164.3 cells/mm) compared to BF controls (1013±178.6 cells/mm, \*\*\**p*=0.0001 vs NEC). In NEC-hPSC animals, enterocytes are increased (882±193.5 cells/mm, \*\**p*=0.005) but do not reach full normal BF levels by Day 4 (\**p*=0.036). (C) OLFM4-positive cells were quantified to assess the presence of LGR5+ ISC within the crypts. As reflected in the reduce crypt height, LGR5+ ISC are significant lost in NEC (19.3±6.4 cells/mm vs BF=61.7±13.5 cells/mm, \*\*\*\**p*<0.0001). With hPSC treatment, LGR5+ ISC re-emerge with a significantly increased population (40.6±10.8 cells/mm, \*\*\*\**p*<0.0001) compared to NEC animals. However, they are not completely restored to BF amounts (\*\**p*=0.0014). (D) While the Paneth cell population is not fully developed in neonates, previous studies have shown their importance in sustaining the ISC niche and in NEC. Larger sections of Lysosyme-immunostained ileum were scanned to more accurately quantify the sparce, developing Paneth cell population. NEC animals had a significant decrease in Paneth cells (8.0±3.0 cells/mm vs BF=14.5±4.3 cells/mm, \*\**p*=0.0046). Paneth cells were significantly increased to slightly above BF levels in NEC animals treated with hPSC (19.1±5.6 cells/mm \*\*\**p*=0.001).

To next assess the intestinal stem cell niche within the crypt, LGR5^+^ intestinal stem cells and Paneth cells were quantified with immunostaining for OLFM4 (a marker of LGR5^+^ stem cells (67)) and Lysozyme (Paneth cells) (Figure 6A, C, D). Compared to BF controls, NEC animals had a significant loss of LGR5^+^ stem cells (BF: 61.7 cells/mm; NEC: 19.3 cells/mm) (Figure 6C). Upon hPSC treatment, the number of the LGR5^+^ stem cells were significantly increased compared to untreated NEC animals (40.6 cells/mm) (Figure 6C). In addition to LGR5^+^ stem cells, Paneth cells are also located within the ileal crypt and play a key role in supporting LGR5^+^ stem cells and in defense (59, 66). Further, recent studies in an experimental NEC mouse model suggest loss of Paneth cells in the presence of aberrant bacterial infection can lead to NEC onset (80). In our experimental NEC rat pup model, Paneth cells are still undergoing expansion and maturation. At this developmental stage, Paneth cells are not always observed within each crypt in 5 µm thick sections of BF control ileum. Instead, they are observed at a rate of 14.5 cells per mm (Figure 6D). As seen in human NEC patients, Paneth cells are significantly lost in NEC pathogenesis with only 8.0 cells per mm in NEC animals (Figure 6D). Similar to LGR5^+^ stem cells, the Paneth cell population was significantly increased to 19.1 cells per mm with hPSC treatment of NEC damage (Figure 6D). These data show that hPSC treatment of NEC results in attenuation of critical epithelial cell losses with a significant re-establishment of the stem cell niche within the crypts.

While LGR5+ stem cells and Paneth cells always reside within the crypts, the distribution of other cellular features may lend insight into the epithelial dynamics in NEC-hPSC animals. Therefore, we examined the cellular distribution of Ezrin-positive enterocytes, EdU-labeled cells, and cleaved caspase 3-positive cells along the crypt-villus axis (Supplemental Figure 2). Localization of LGR5+ stem cells was used as a localization control as their cellular position remains mostly unchanged within the crypt. First, localization of EdU-positive epithelial cells was assessed to examine potential cellular migration of the recently proliferating cells. Histograms of EdU-positive cell distribution shows proliferating cells in BF animals migrate out of the crypts towards the villus tip with occasional EdU-positive cells found further up the villi (Supplemental Figure 2). In NEC animals, EdU-positive cells appear to have decreased migration from the crypts. Upon hPSC treatment, the number of EdU-positive cells increase (Figure 5F). However, most of these cells are localized near the crypts with relatively no EdU-positive cells observed up the villi as observed in BF controls (Supplemental Figure 2). In contrast, apoptotic epithelial cells were more widely dispersed along the crypt-villus axis (Supplemental Figure 2). One increased region of apoptotic cells within the villi was noted in NEC animals. Interestingly, when comparing enterocyte distributions between NEC and NEC-hPSC animals, NEC animals displayed a decrease in enterocytes in the same general villar localization as the aberrant apoptosis. This may suggest that increased apoptosis within this villar region corresponds with the decrease in enterocytes and resultant shortened villar height. As Edu-positive cells do not rapidly migrate up the villi in NEC-hPSC animals, the increase in enterocytes beginning at this same localization may be the result of the observed resolution of this aberrant apoptosis rather than simply replenishment via newly divided cells. As expected, LGR5+ stem cells and Paneth cells (not shown) showed increased numbers of cells but no changes in distribution between animal groups with localization consistently within the crypts. Together, this in-depth distribution analysis yields insight into the complex interplay of cellular changes that occur in enterocytes along the villi in addition to the intestinal stem cell niche depletion and expansion. Further, this method provides an additional tool for future analysis of the complex cellular dynamics involved in cessation and repair of NEC damage.

## Discussion

In the last decade, researchers have made significant advances in identifying important prophylactic strategies for reducing the risk of NEC onset (54, 64). Unfortunately, few approaches have demonstrated the therapeutic ability to ameliorate established NEC damage. To address this gap, we focused our present study on the potential of hPSC for NEC treatment. In the neonatal rat model of NEC, intestinal damage is induced within 24-36 hours from the start of the NEC stressors (6, 58, 79) and confirmed through observations of clinical symptoms onset shown in Figure 2. Due to increasing mortality rates, this model is limited to a 96-hour endpoint. Based on these timings, our present study timeline included hPSC injection following the conclusion of LPS administration. While this resulted in a shortened treatment timetable, it did enable investigation of therapeutic benefits as NEC intestinal damage was induced prior to treatment. An additional challenge for NEC therapeutic studies is clinical application, specifically pertaining to the route of administration. Intravenous administration of cell therapies is associated with cell entrapment and embolism risks in the lungs (20, 74). Therefore, we sought an alternative administration route to avoid these potential risk factors in premature infants already predisposed to lung impairment. Previous studies have supported the safety and efficacy of intraperitoneal (IP) injection for cell therapy including preclinical NEC investigations (23, 68, 73). Considering these clinical translation factors, the IP route of administration was chosen to balance the potential risks and benefits in this unique patient population.

In this study, we examined the therapeutic effects across the multiple scales ranging from whole animal clinical symptoms to histological cellular damage using comprehensive quantitative analyses at each length scale. At the cellular level, the villi/crypt morphologies and essential epithelial cell populations were re-established in hPSC treated NEC pups. As assessed by the ‘NEC Grade’ composite score, the overall severity of histological damage across the ileum was significantly decreased. Additionally, hPSC therapy also reduced the amount of ileal tissue with NEC damage. This cellular level amelioration extended to observed improvement at the macroscopic level. Upon gross examination, the ‘Macroscopic Score’ that included fragility and distention was decreased in hPSC-NEC ileum. Similar to the histological assessment, the extent of ileal tissue involved in the pathological damage was also significantly reduced. However, these cellular and tissue level recoveries did not correlate to full resolution of whole animal clinical symptoms. Although trending towards improvement, body weight and Clinical Sickness Score did not reach statistical significance by the 96-hour endpoint. A previous study using amniotic fluid stem cells for NEC treatment has shown whole animal improvements with cells administered at 24 hours (77, 78). Our current study was designed to allow NEC damage to be further initiated and established (6, 58, 79) prior to our 32-hour treatment timepoint (8-hour increase in NEC initiation time). The resultant shortened treatment window could be a contributing reason for the lack of statistically significant clinical resolution by the study endpoint. We also specifically utilized human-derived PSC to maintain clinical relevance, however, potential decreased efficacy due to the species mismatch should also be considered. Additionally, other non-intestinal organs have been shown to incur damage and contribute to advancing NEC pathogenesis and long-term adversities (9, 29, 53). Future in-depth studies of hPSC localization not investigated here and potential ‘extra-intestinal’ effects of hPSC therapy may lend insight into potential therapeutic benefits for non-intestinal damage in addition to addressing the gap between observed histological/intestinal level improvement and lack of whole animal clinical symptom resolution.

NEC is characteristically a multi-focal disease in the premature ileum. With current medical management and supportive care during early stage NEC, over 30% of patients progress to advanced disease requiring surgical intervention to remove necrotic segments (17). This unpredictable progression can occur within days or even hours and correlates with drastically increased mortality rates (up to 50%) (21, 22). For survivors, surgical management is also compounded by long term complications associated to the amount of remaining healthy bowel remaining (14, 16, 31, 57). Therefore, both the severity and the amount of tissue involved in NEC are important factors in assessing potential therapeutic approaches. To address this, we quantified the overall severity and the extent of tissue involvement for each of the NEC Grades. hPSC therapy decreased the amount of ileal tissue involved in pathogenic NEC damage from 25% to less than 5%. Moreover, an NEC Grade of 3 or above was only rarely observed in hPSC-NEC animals. If translated into a clinical setting, this could result in a smaller portion of bowel removed or possible avoidance of surgery all together for some patients. The reduction of surgical interventions alone could significantly impact outcomes for NEC patients. In addition to this histological assessment, we also measured the severity and extent of the macroscopic NEC pathology. As seen from the breakdown of the composite ‘Macroscopic Score’, intestinal distention was a key factor in the overall pathology score. In humans, pathologically distended intestine can cause impaired motility and predispose the intestine to dysbiosis and malabsorption (10, 12, 33), and thus we sought to further investigate the severity and extent of the ileal dilation. As the severe NEC distention was observed in a multi-focal pattern along the length of the ileum, a custom software was developed to automate the quantification of both the severity of distention and the percent of ileal tissue involved in this ‘patchy’ pathology. NEC was shown to cause significant distention pathology in almost 25% of the ileum. hPSC therapy reduces the amount of tissue involved to less than 10%. A comparison of gross pathology to the histological damage analysis also revealed a similar percentage of tissue involved in each. This finding supports a previously reported correlation between the severity of the ‘Macroscopic Score’ as an indicator of the ‘NEC Grade’ (79). While previously used composite scores can give an overall assessment of the tissue, more in-depth quantitative analysis of tissue involvement can be beneficial when investigating a multi-focal disease such as NEC. This newly developed software for automated pathological distention measurements now provides a novel research analysis tool that reduces site selection or perception bias, increases throughput, and allows for more complete multi-level comparisons of NEC damage in future studies.

As NEC is a multifactorial disease involving several instigating factors, multiple potential therapeutic targets have been theorized. Of these, reduction of the over-reactive immune response is a common focus of therapeutic strategies as it plays a significant role in not only intestinal NEC damage but also systemic sepsis. However, clinical application is a challenge for NEC therapeutics that target narrow suppression of the pathogenic inflammation. For example, inhibitors that suppress the specific TNFα cytokine in the inflammatory milieu have shown therapeutic ability in preclinical NEC models (65), but also pose an uncertain risk of systemic side effects (i.e. further infections) in an already vulnerable patient population (18). Perinatal stem cells present a unique approach known for broader immune modulation rather than specific inflammatory suppression (36, 62). hPSC also uniquely secrete a multitude of factors that can target other components of NEC pathogenesis via potent paracrine signaling (27, 37, 41). Therefore, we hypothesized that hPSC therapy was a novel multi-faceted paracrine approach that could improve epithelial damage and reduce pathogenic inflammation in NEC disease. As discussed above, hPSC therapy improved macroscopic and histological outcomes in NEC disease as the result of potentially two synergistic sources: prevention of disease progression and promotion of epithelial recovery. To examine changes in the pathogenic inflammation, we used a panel of the 3 prominent inflammatory biomarkers involved in NEC pathogenesis: TNFα, IL1β, and NFκB (4, 58, 61, 72, 76). In the ileum, we found hPSC therapy significantly reduced TNFα levels. However, IL1β and NFκB levels did not reach statistical significance in their decreasing expression trend. While we expected broad reduction in pathogenic inflammation, these data suggest only a moderate decrease across the 3 inflammatory signaling biomarkers upon hPSC therapy. Interestingly, the observed hPSC-induced reduction of TNFα does support the previous TNFα inhibitor study’s findings that TNFα plays an important role in NEC pathogenesis in the intestine (61). However, the moderate reduction in pro-inflammatory signaling markers suggest hPSC do not induce NEC healing through potent reduction of pathogenic inflammation. Conversely, recent studies have shown an increase in specific reparative immune pathways during NEC recovery in patients (52, 72) and future studies could investigate these specific reparative immune pathways not examined in the present study. While NEC is traditionally thought of as a heavily immune-driven disease, these data with only moderate reduction in pathogenic inflammation suggest that support of epithelial repair can also be an effective target in NEC treatment approaches.

One of the most prominent findings in our present study is the significant improvement of the intestinal stem cell niche, specifically the LGR5+ stem cell and Paneth cell populations. Recent studies have begun to highlight the importance of these two epithelial cell lineages in NEC. In *in vitro* studies with NEC enteroids, stimulation of LGR5+ stem cells with exogenous Wnt3a was shown to promote epithelial survival and growth (38). Without exogenous Wnt3a administration to stimulate LGR5+ cells, fewer NEC enteroids survived in culture. Moreover, Paneth cells are endogenous sources of Wnt signaling that supports LGR5+ stem cells in healthy crypts. Loss of Paneth cells coupled with bacterial colonization in neonate mice lead to the onset of characteristic NEC disease (80). These investigations have begun to reveal the importance of both Paneth cells and LGR5+ stem cells in prevention and recovery of NEC damage. In our NEC animal model, the loss of Paneth cells and LGR5+ stem cells observed in NEC was ameliorated upon hPSC treatment. These findings coupled with the mild reduction in pathogenic inflammation suggest the stem cell niche support is a key mechanism of action utilized by hPSC therapy.

In intestinal injuries associated with stem cell niche loss, several signaling pathways have been identified to support the subsequent repair of the niche. In wound healing of ulcerated lesions, localization of Cox2-expressing cells abutting the crypt is essential for epithelial repair and replenishment (46). A role for crypt-adjacent stromal Cox2+ cells has also been suggested in amniotic fluid stem cell treatment of experimental NEC (77). Furthermore, administration of a Cox2 inhibitor negated the reparative effects of amniotic fluid stem cell therapy on the survival outcome of NEC animals (77). Thus, a similar pathway may also be utilized by hPSC therapy to repair the intestinal stem cell niche with NEC damage. In addition, hPSC also have been recently shown to uniquely express EGF receptor (EGFR) and NRG1, members of the EGFR/ErbB signaling pathways (41). Activation of EGFR via breastmilk exposure is a prominent protective pathway in the neonatal intestinal epithelium. Notably, exogenous dosing of EGF and HB-EGF, EGFR activating ligands, can protect against NEC onset but fail to repair established experimental NEC damage (56, 73). The therapeutic insufficiency of exogenous ligand dosing may be due to the significant reduction of EGFR expression that occurs in NEC pathogenesis (25). However, previous studies have demonstrated the ability of secreted EGFR-containing exosomes to increase EGFR expression and possibly amplify downstream signaling in recipient cells (30, 81). Further studies are needed to investigate if this can occur with hPSC-secreted EGFR into NEC damaged tissue, specifically LGR5+ stem cells. In a similar signaling pathway, hPSC also express NRG1 (41), a ligand for the Erb4 receptor in the EGFR/Erb receptor kinase family. In cultured mouse ileal enteroids, Erb4 loss via genetic deletion reduces the Paneth cell population. Further activation of Erb4 in an experimental NEC model protected the Paneth cell lineage. As we observed significant expansion of Paneth cells to even higher than BF levels, hPSC may also signal through the EGFR/ErbB family to protect or increase Paneth cell lineage that can in turn support the LGR5+ stem cells and combat the ongoing bacterial translocation. In addition to these more well-known pathways in NEC, several other factors have also been found in the hPSC secretome such as Lactoferrin, ICAM-1, EpCam, and Integrin β1 that can target intestinal stem cells in homeostasis and other intestinal disease (41, 55). Altogether, hPSC possess a plethora of potential signaling pathways including both direct to epithelium signaling (EGFR/ErbB4) and indirect cellular interactions (Cox2+ stromal cells) that could be potential mechanisms of action that critically support the stem cell niche restoration observed in our current study. While future studies can further elucidate the pivotal hPSC signaling pathways, the findings here establish that support of the intestinal stem cell niche is a key mechanism of hPSC treatment for NEC disease.

In conclusion, hPSC therapy can ameliorate established NEC damage at the cellular and tissue level in the NEC animal model. While broad reduction in pathogenic inflammatory cytokines was not observed, significant epithelial and stem cell niche amelioration corresponding to improved macroscopic health was induced with hPSC treatment. As perinatal stem cell therapies have shown low long-term engraftment rates into the intestine (77), it is reasonable to assume hPSC therapy functions via paracrine signaling similar to that observed in previous studies (41, 77). In phase I clinical trials, perinatal stem cells appear to have an acceptable safety profile for use in premature infants (1, 2, 8). Therefore, these findings demonstrate hPSC are a unique approach that should be further investigated for clinical translation as a NEC therapeutic strategy.

Future studies of hPSC-based therapy will include dose range studies, in-depth investigations into potential reparative immunomodulation, and hPSC secretome administration (i.e. conditioned media, exosomes) to conceivably increase safety and efficacy. Additionally, hPSC also provide a critical research tool that can be utilized to elucidate the necessary mechanisms in prevention of progression and repair to expand our overall understanding of neonatal repair pathophysiology.

---

**Supplemental Figure 1.**
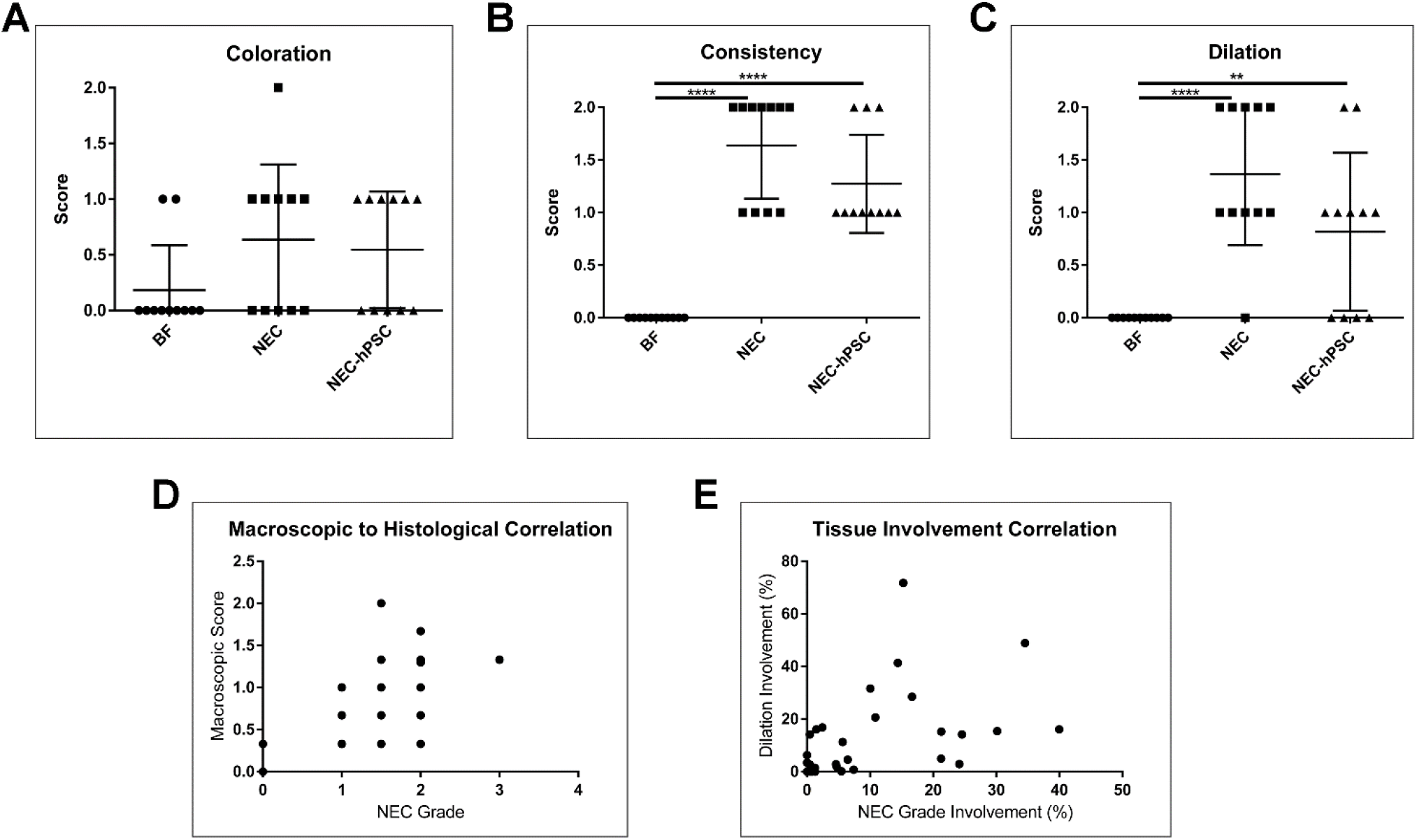
Gross pathology assessment and histological correlation. The composite Macroscopic Score used to assess the gross pathological health of the intestine is composed of 3 factors: Coloration, Consistency, and Dilation. In panels A, B, and C, the breakdown of the individual factors is plotted. Coloration has little effect on the increase in the Macroscopic Score observed in NEC animals. While consistency and dilation both contribute to the increased NEC Macroscopic Score compared to NEC \*\*\*\**p*<0.0001), dilation appears to be a key factor that drives the amelioration of the gross pathological damage in NEC-hPSC closer to BF levels (\*\**p*=0.0039, NEC-hPSC vs BF). Based on these findings, more in-depth analysis was performed in this study. (D) As previously reported, the gross pathological Macroscopic Score shows correlation to the histological NEC Grade of each animal (r=0.778, *p*<0.0001). (E) As dilation is a score driving factor of the gross pathological damage, the correlation of the amount of tissue with this pathogenic dilation to the amount of tissue with histological damage (NEC Grade ≥2) was assessed. This correlation analysis suggests there may be a correlation between tissue involvement at the histological and the macroscopic level (r=0.6564, *p*<0.0001). Future studies could further investigate the utility of this correlation as an additional research tool for in-depth assessment of multi-scale NEC damage.

**Supplemental Figure 2.**
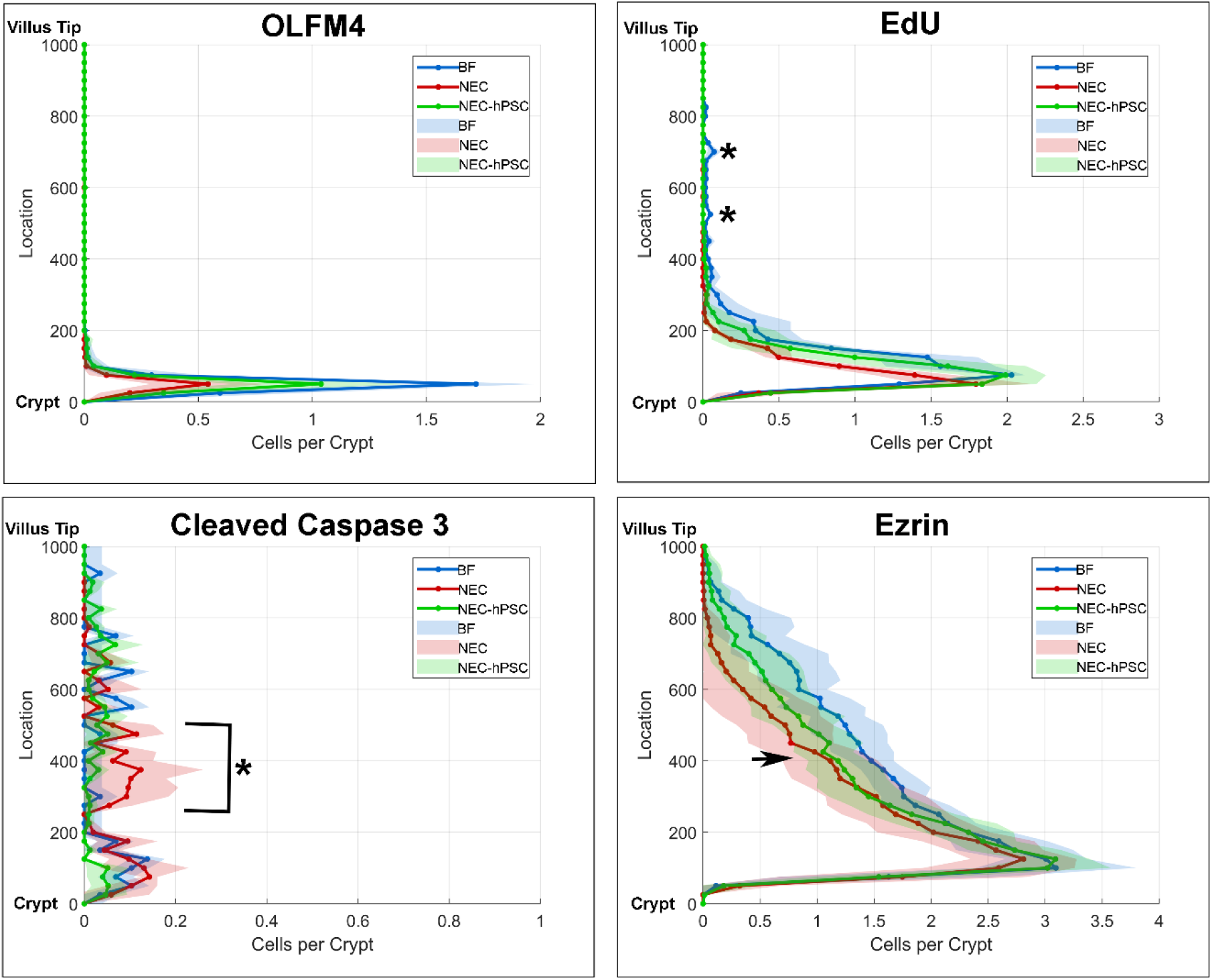
Assessment of cellular distribution. Using the same immunostained sections as for previous quantifications, the localization of each labeled cell was measured as the distance from the crypt base. The histogram depicts the average number of cells per crypt for each location within the crypt-villus axis as depicted on the Y-axis (0 pixels=crypt base, increasing location/pixel number on Y-axis signifies increasing distance away from crypt base). The colored shading represents the standard deviation for each point along the histogram. Besides an increase in the total number of OLFM4-positive LGR5+ stem cells (i.e. area under the curve), no changes were observed in the distribution of intestinal stem cells between the animal groups (top left panel). EdU-labeled proliferating cells were occasionally observed in the upper portions of the villi (asterisk, top right panel). Interestingly, the increased proliferation found in NEC-hPSC animals did not show significant localization up the villi but appeared to be more localized to the crypt region where an expansion of LGR5+ stem cells and Paneth cells have been found. An aberrant pattern of apoptotic epithelial cells was noted in NEC animals (bracketed asterisk, bottom left panel). This pattern was localized above the crypt region and corresponded to the differentiating cells of the villi. The increased enterocyte numbers found in Figure 6 were localized a greater distance from the crypt than NEC enterocytes (bottom right panel). Further, the localization of the increased NEC-hPSC enterocytes diverged near the same region as the aberrant apoptotic pattern (arrow). These cellular distributions provide insight into the cellular damage and healing involved in NEC and hPSC treatment.

## Grants

These studies were supported by the state of North Carolina (A.A), Lisa Dean-Moseley Foundation (M.Z.S. and V.G.W), and NIH-NCI K25CA204599 (J.A.W.).

## Disclosures

## Acknowledgments

The Ezrin antibody developed by Clinical Proteomics Technologies for Cancer was obtained from the Developmental Studies Hybridoma Bank, created by the NICHD of the NIH and maintained at The University of Iowa, Department of Biology, Iowa City, IA. We thank Dr. Frank Marini and Kristina Stumpf for their assistance with the Nuance system for immunofluorescent imaging. We also thank the Regenerative Medicine Clinical Center at Wake Forest Institute for Regenerative Medicine for providing the human placental-derived stem cells.

## Notes

### Competing Interest Statement

The authors have declared no competing interest.

## References

1. Ahn SY, Chang YS, Kim JH, Sung SI, and Park WS. Two-Year Follow-Up Outcomes of Premature Infants Enrolled in the Phase I Trial of Mesenchymal Stem Cells Transplantation for Bronchopulmonary Dysplasia. J Pediatr 185: 49–54 e42, 2017.

2. Ahn SY, Chang YS, Sung SI, and Park WS. Mesenchymal Stem Cells for Severe Intraventricular Hemorrhage in Preterm Infants: Phase I Dose-Escalation Clinical Trial. Stem Cells Transl Med 7: 847–856, 2018.

3. Akira S, Takeda K, and Kaisho T. Toll-like receptors: critical proteins linking innate and acquired immunity. Nat Immunol 2: 675–680, 2001.

4. Baregamian N, Song J, Bailey CE, Papaconstantinou J, Evers BM, and Chung DH. Tumor necrosis factor-alpha and apoptosis signal-regulating kinase 1 control reactive oxygen species release, mitochondrial autophagy, and c-Jun N-terminal kinase/p38 phosphorylation during necrotizing enterocolitis. Oxid Med Cell Longev 2: 297–306, 2009.

5. Barlow B, Santulli TV, Heird WC, Pitt J, Blanc WA, and Schullinger JN. An experimental study of acute neonatal enterocolitis--the importance of breast milk. J Pediatr Surg 9: 587–595, 1974.

6. Bergmann KR, Liu SX, Tian R, Kushnir A, Turner JR, Li HL, Chou PM, Weber CR, and De Plaen IG. Bifidobacteria stabilize claudins at tight junctions and prevent intestinal barrier dysfunction in mouse necrotizing enterocolitis. Am J Pathol 182: 1595–1606, 2013.

7. Cetin S, Ford HR, Sysko LR, Agarwal C, Wang J, Neal MD, Baty C, Apodaca G, and Hackam DJ. Endotoxin inhibits intestinal epithelial restitution through activation of Rho-GTPase and increased focal adhesions. J Biol Chem 279: 24592–24600, 2004.

8. Chang YS, Ahn SY, Yoo HS, Sung SI, Choi SJ, Oh WI, and Park WS. Mesenchymal stem cells for bronchopulmonary dysplasia: phase 1 dose-escalation clinical trial. J Pediatr 164: 966–972 e966, 2014.

9. Criss CN, Selewski DT, Sunkara B, Gish JS, Hsieh L, McLeod JS, Robertson JO, Matusko N, and Gadepalli SK. Acute kidney injury in necrotizing enterocolitis predicts mortality. Pediatr Nephrol 33: 503–510, 2018.

10. D’Antiga L, and Goulet O. Intestinal failure in children: the European view. J Pediatr Gastroenterol Nutr 56: 118–126, 2013.

11. De Coppi P, Bartsch G, Jr., Siddiqui MM, Xu T, Santos CC, Perin L, Mostoslavsky G, Serre AC, Snyder EY, Yoo JJ, Furth ME, Soker S, and Atala A. Isolation of amniotic stem cell lines with potential for therapy. Nat Biotechnol 25: 100–106, 2007.

12. Dibaise JK, Young RJ, and Vanderhoof JA. Enteric microbial flora, bacterial overgrowth, and short-bowel syndrome. Clin Gastroenterol Hepatol 4: 11–20, 2006.

13. Drucker NA, McCulloh CJ, Li B, Pierro A, Besner GE, and Markel TA. Stem cell therapy in necrotizing enterocolitis: Current state and future directions. Semin Pediatr Surg 27: 57–64, 2018.

14. Duro D, Kalish LA, Johnston P, Jaksic T, McCarthy M, Martin C, Dunn JC, Brandt M, Nobuhara KK, Sylvester KG, Moss RL, and Duggan C. Risk factors for intestinal failure in infants with necrotizing enterocolitis: a Glaser Pediatric Research Network study. J Pediatr 157: 203–208 e201, 2010.

15. Egan CE, Sodhi CP, Good M, Lin J, Jia H, Yamaguchi Y, Lu P, Ma C, Branca MF, Weyandt S, Fulton WB, Nino DF, Prindle T, Jr., Ozolek JA, and Hackam DJ. Toll-like receptor 4-mediated lymphocyte influx induces neonatal necrotizing enterocolitis. J Clin Invest 126: 495–508, 2016.

16. Elfvin A, Dinsdale E, Wales PW, and Moore AM. Low birthweight, gestational age, need for surgical intervention and gram-negative bacteraemia predict intestinal failure following necrotising enterocolitis. Acta Paediatr 104: 771–776, 2015.

17. Ellsbury DL, Clark RH, Ursprung R, Handler DL, Dodd ED, and Spitzer AR. A Multifaceted Approach to Improving Outcomes in the NICU: The Pediatrix 100 000 Babies Campaign. Pediatrics 137: 2016.

18. Esteve-Sole A, Deya-Martinez A, Teixido I, Ricart E, Gompertz M, Torradeflot M, de Moner N, Gonzalez EA, Plaza-Martin AM, Yague J, Juan M, and Alsina L. Immunological Changes in Blood of Newborns Exposed to Anti-TNF-alpha during Pregnancy. Front Immunol 8: 1123, 2017.

19. Fanaroff AA, Stoll BJ, Wright LL, Carlo WA, Ehrenkranz RA, Stark AR, Bauer CR, Donovan EF, Korones SB, Laptook AR, Lemons JA, Oh W, Papile LA, Shankaran S, Stevenson DK, Tyson JE, Poole WK, and Network NNR. Trends in neonatal morbidity and mortality for very low birthweight infants. Am J Obstet Gynecol 196: 147 e141–148, 2007.

20. Fischer UM, Harting MT, Jimenez F, Monzon-Posadas WO, Xue H, Savitz SI, Laine GA, and Cox CS, Jr. Pulmonary passage is a major obstacle for intravenous stem cell delivery: the pulmonary first-pass effect. Stem Cells Dev 18: 683–692, 2009.

21. Fitzgibbons SC, Ching Y, Yu D, Carpenter J, Kenny M, Weldon C, Lillehei C, Valim C, Horbar JD, and Jaksic T. Mortality of necrotizing enterocolitis expressed by birth weight categories. J Pediatr Surg 44: 1072-1075; discussion 1075-1076, 2009.

22. Fredriksson F, and Engstrand Lilja H. Survival rates for surgically treated necrotising enterocolitis have improved over the last four decades. Acta Paediatr 108: 1603–1608, 2019.

23. Ghionzoli M, Cananzi M, Zani A, Rossi CA, Leon FF, Pierro A, Eaton S, and De Coppi P. Amniotic fluid stem cell migration after intraperitoneal injection in pup rats: implication for therapy. Pediatr Surg Int 26: 79–84, 2010.

24. Goncalves FL, Gallindo RM, Soares LM, Figueira RL, Volpe FA, Pereira-da-Silva MA, and Sbragia L. Validation of protocol of experimental necrotizing enterocolitis in rats and the pitfalls during the procedure. Acta Cir Bras 028 Suppl 1: 19–25, 2013.

25. Good M, Siggers RH, Sodhi CP, Afrazi A, Alkhudari F, Egan CE, Neal MD, Yazji I, Jia H, Lin J, Branca MF, Ma C, Prindle T, Grant Z, Shah S, Slagle D, 2nd, Paredes J, Ozolek J, Gittes GK, and Hackam DJ. Amniotic fluid inhibits Toll-like receptor 4 signaling in the fetal and neonatal intestinal epithelium. Proc Natl Acad Sci U S A 109: 11330–11335, 2012.

26. Good M, Sodhi CP, Egan CE, Afrazi A, Jia H, Yamaguchi Y, Lu P, Branca MF, Ma C, Prindle T, Jr., Mielo S, Pompa A, Hodzic Z, Ozolek JA, and Hackam DJ. Breast milk protects against the development of necrotizing enterocolitis through inhibition of Toll-like receptor 4 in the intestinal epithelium via activation of the epidermal growth factor receptor. Mucosal Immunol 8: 1166–1179, 2015.

27. Guan YT, Xie Y, Li DS, Zhu YY, Zhang XL, Feng YL, Chen YP, Xu LJ, Liao PF, and Wang G. Comparison of biological characteristics of mesenchymal stem cells derived from the human umbilical cord and decidua parietalis. Mol Med Rep 20: 633–639, 2019.

28. Gurien LA, Stallings-Archer K, and Smith SD. Probiotic Lactococcus lactis decreases incidence and severity of necrotizing enterocolitis in a preterm animal model. J Neonatal Perinatal Med 11: 65–69, 2018.

29. Halpern MD, Holubec H, Dominguez JA, Meza YG, Williams CS, Ruth MC, McCuskey RS, and Dvorak B. Hepatic inflammatory mediators contribute to intestinal damage in necrotizing enterocolitis. Am J Physiol Gastrointest Liver Physiol 284: G695–702, 2003.

30. Higginbotham JN, Zhang Q, Jeppesen DK, Scott AM, Manning HC, Ochieng J, Franklin JL, and Coffey RJ. Identification and characterization of EGF receptor in individual exosomes by fluorescence-activated vesicle sorting. J Extracell Vesicles 5: 29254, 2016.

31. Hintz SR, Kendrick DE, Stoll BJ, Vohr BR, Fanaroff AA, Donovan EF, Poole WK, Blakely ML, Wright L, Higgins R, and Network NNR. Neurodevelopmental and growth outcomes of extremely low birth weight infants after necrotizing enterocolitis. Pediatrics 115: 696–703, 2005.

32. Horbar JD, Carpenter JH, Badger GJ, Kenny MJ, Soll RF, Morrow KA, and Buzas JS. Mortality and neonatal morbidity among infants 501 to 1500 grams from 2000 to 2009. Pediatrics 129: 1019–1026, 2012.

33. Hukkinen M, Mutanen A, and Pakarinen MP. Small bowel dilation in children with short bowel syndrome is associated with mucosal damage, bowel-derived bloodstream infections, and hepatic injury. Surgery 162: 670–679, 2017.

34. Ip S, Chung M, Raman G, Chew P, Magula N, DeVine D, Trikalinos T, and Lau J. Breastfeeding and maternal and infant health outcomes in developed countries. Evid Rep Technol Assess (Full Rep) 1–186, 2007.

35. Jilling T, Lu J, Jackson M, and Caplan MS. Intestinal epithelial apoptosis initiates gross bowel necrosis in an experimental rat model of neonatal necrotizing enterocolitis. Pediatr Res 55: 622–629, 2004.

36. Khoury O, Atala A, and Murphy SV. Stromal cells from perinatal and adult sources modulate the inflammatory immune response in vitro by decreasing Th1 cell proliferation and cytokine secretion. Stem Cells Transl Med 2019.

37. Kong P, Xie X, Li F, Liu Y, and Lu Y. Placenta mesenchymal stem cell accelerates wound healing by enhancing angiogenesis in diabetic Goto-Kakizaki (GK) rats. Biochem Biophys Res Commun 438: 410–419, 2013.

38. Li B, Lee C, Cadete M, Zhu H, Koike Y, Hock A, Wu RY, Botts SR, Minich A, Alganabi M, Chi L, Zani-Ruttenstock E, Miyake H, Chen Y, Mutanen A, Ngan B, Johnson-Henry KC, De Coppi P, Eaton S, Maattanen P, Delgado-Olguin P, Sherman PM, Zani A, and Pierro A. Impaired Wnt/beta-catenin pathway leads to dysfunction of intestinal regeneration during necrotizing enterocolitis. Cell Death Dis 10: 743, 2019.

39. Maheshwari A, Kelly DR, Nicola T, Ambalavanan N, Jain SK, Murphy-Ullrich J, Athar M, Shimamura M, Bhandari V, Aprahamian C, Dimmitt RA, Serra R, and Ohls RK. TGF-beta2 suppresses macrophage cytokine production and mucosal inflammatory responses in the developing intestine. Gastroenterology 140: 242–253, 2011.

40. Mareschi K, Castiglia S, Sanavio F, Rustichelli D, Muraro M, Defedele D, Bergallo M, and Fagioli F. Immunoregulatory effects on T lymphocytes by human mesenchymal stromal cells isolated from bone marrow, amniotic fluid, and placenta. Exp Hematol 44: 138–150 e131, 2016.

41. Matz EL, Thakker PU, Gu X, Terlecki RP, Dou L, Walker SJ, Lue T, Lin G, Atala A, Yoo JJ, Zhang Y, and Jackson JD. Administration of secretome from human placental stem cell-conditioned media improves recovery of erectile function in the pelvic neurovascular injury model. J Tissue Eng Regen Med 2020.

42. Maurer CR, Qi R, and Raghavan V. A linear time algorithm for computing exact Euclidean distance transforms of binary images in arbitrary dimensions. IEEE Transactions on Pattern Analysis and Machine Intelligence 25: 265–270, 2003.

43. McCulloh CJ, Olson JK, Wang Y, Zhou Y, Tengberg NH, Deshpande S, and Besner GE. Treatment of experimental necrotizing enterocolitis with stem cell-derived exosomes. J Pediatr Surg 53: 1215–1220, 2018.

44. McCulloh CJ, Olson JK, Zhou Y, Wang Y, and Besner GE. Stem cells and necrotizing enterocolitis: A direct comparison of the efficacy of multiple types of stem cells. J Pediatr Surg 52: 999–1005, 2017.

45. McElroy SJ, Underwood MA, and Sherman MP. Paneth cells and necrotizing enterocolitis: a novel hypothesis for disease pathogenesis. Neonatology 103: 10–20, 2013.

46. Miyoshi H, VanDussen KL, Malvin NP, Ryu SH, Wang Y, Sonnek NM, Lai CW, and Stappenbeck TS. Prostaglandin E2 promotes intestinal repair through an adaptive cellular response of the epithelium. EMBO J 36: 5–24, 2017.

47. MohanKumar K, Kaza N, Jagadeeswaran R, Garzon SA, Bansal A, Kurundkar AR, Namachivayam K, Remon JI, Bandepalli CR, Feng X, Weitkamp JH, and Maheshwari A. Gut mucosal injury in neonates is marked by macrophage infiltration in contrast to pleomorphic infiltrates in adult: evidence from an animal model. Am J Physiol Gastrointest Liver Physiol 303: G93–102, 2012.

48. MohanKumar K, Namachivayam K, Chapalamadugu KC, Garzon SA, Premkumar MH, Tipparaju SM, and Maheshwari A. Smad7 interrupts TGF-beta signaling in intestinal macrophages and promotes inflammatory activation of these cells during necrotizing enterocolitis. Pediatr Res 79: 951–961, 2016.

49. Moorefield EC, McKee EE, Solchaga L, Orlando G, Yoo JJ, Walker S, Furth ME, and Bishop CE. Cloned, CD117 selected human amniotic fluid stem cells are capable of modulating the immune response. PLoS One 6: e26535, 2011.

50. Morgan RL, Preidis GA, Kashyap PC, Weizman AV, Sadeghirad B, McMaster Probiotic P, and Synbiotic Work G. Probiotics Reduce Mortality and Morbidity in Preterm, Low-Birth-Weight Infants: A Systematic Review and Network Meta-analysis of Randomized Trials. Gastroenterology 2020.

51. Neal MD, Sodhi CP, Jia H, Dyer M, Egan CE, Yazji I, Good M, Afrazi A, Marino R, Slagle D, Ma C, Branca MF, Prindle T, Jr., Grant Z, Ozolek J, and Hackam DJ. Toll-like receptor 4 is expressed on intestinal stem cells and regulates their proliferation and apoptosis via the p53 up-regulated modulator of apoptosis. J Biol Chem 287: 37296–37308, 2012.

52. Nino DF, Sodhi CP, Egan CE, Zhou Q, Lin J, Lu P, Yamaguchi Y, Jia H, Martin LY, Good M, Fulton WB, Prindle T, Jr., Ozolek JA, and Hackam DJ. Retinoic Acid Improves Incidence and Severity of Necrotizing Enterocolitis by Lymphocyte Balance Restitution and Repopulation of LGR5+ Intestinal Stem Cells. Shock 47: 22–32, 2017.

53. Nino DF, Zhou Q, Yamaguchi Y, Martin LY, Wang S, Fulton WB, Jia H, Lu P, Prindle T, Jr., Zhang F, Crawford J, Hou Z, Mori S, Chen LL, Guajardo A, Fatemi A, Pletnikov M, Kannan RM, Kannan S, Sodhi CP, and Hackam DJ. Cognitive impairments induced by necrotizing enterocolitis can be prevented by inhibiting microglial activation in mouse brain. Sci Transl Med 10: 2018.

54. Patel RM, and Underwood MA. Probiotics and necrotizing enterocolitis. Semin Pediatr Surg 27: 39–46, 2018.

55. Peak TC, Praharaj PP, Panigrahi GK, Doyle M, Su Y, Schlaepfer IR, Singh R, Vander Griend DJ, Alickson J, Hemal A, Atala A, and Deep G. Exosomes secreted by placental stem cells selectively inhibit growth of aggressive prostate cancer cells. Biochem Biophys Res Commun 499: 1004–1010, 2018.

56. Radulescu A, Zorko NA, Yu X, and Besner GE. Preclinical neonatal rat studies of heparin-binding EGF-like growth factor in protection of the intestines from necrotizing enterocolitis. Pediatr Res 65: 437–442, 2009.

57. Rees CM, Pierro A, and Eaton S. Neurodevelopmental outcomes of neonates with medically and surgically treated necrotizing enterocolitis. Arch Dis Child Fetal Neonatal Ed 92: F193–198, 2007.

58. Rentea RM, Welak SR, Fredrich K, Donohoe D, Pritchard KA, Oldham KT, Gourlay DM, and Liedel JL. Early enteral stressors in newborns increase inflammatory cytokine expression in a neonatal necrotizing enterocolitis rat model. Eur J Pediatr Surg 23: 39–47, 2013.

59. Sato T, van Es JH, Snippert HJ, Stange DE, Vries RG, van den Born M, Barker N, Shroyer NF, van de Wetering M, and Clevers H. Paneth cells constitute the niche for Lgr5 stem cells in intestinal crypts. Nature 469: 415–418, 2011.

60. Schaart MW, de Bruijn AC, Bouwman DM, de Krijger RR, van Goudoever JB, Tibboel D, and Renes IB. Epithelial functions of the residual bowel after surgery for necrotising enterocolitis in human infants. J Pediatr Gastroenterol Nutr 49: 31–41, 2009.

61. Seitz G, Warmann SW, Guglielmetti A, Heitmann H, Ruck P, Kreis ME, and Fuchs J. Protective effect of tumor necrosis factor alpha antibody on experimental necrotizing enterocolitis in the rat. J Pediatr Surg 40: 1440–1445, 2005.

62. Silini AR, Magatti M, Cargnoni A, and Parolini O. Is Immune Modulation the Mechanism Underlying the Beneficial Effects of Amniotic Cells and Their Derivatives in Regenerative Medicine? Cell Transplant 26: 531–539, 2017.

63. Smart JL, Massey RF, Nash SC, and Tonkiss J. Effects of early-life undernutrition in artificially reared rats: subsequent body and organ growth. Br J Nutr 58: 245–255, 1987.

64. Sullivan S, Schanler RJ, Kim JH, Patel AL, Trawoger R, Kiechl-Kohlendorfer U, Chan GM, Blanco CL, Abrams S, Cotten CM, Laroia N, Ehrenkranz RA, Dudell G, Cristofalo EA, Meier P, Lee ML, Rechtman DJ, and Lucas A. An exclusively human milk-based diet is associated with a lower rate of necrotizing enterocolitis than a diet of human milk and bovine milk-based products. J Pediatr 156: 562–567 e561, 2010.

65. Tayman C, Aydemir S, Yakut I, Serkant U, Ciftci A, Arslan E, and Koc O. TNF-alpha Blockade Efficiently Reduced Severe Intestinal Damage in Necrotizing Enterocolitis. J Invest Surg 29: 209–217, 2016.

66. Vaishnava S, Behrendt CL, Ismail AS, Eckmann L, and Hooper LV. Paneth cells directly sense gut commensals and maintain homeostasis at the intestinal host-microbial interface. Proc Natl Acad Sci U S A 105: 20858–20863, 2008.

67. van der Flier LG, Haegebarth A, Stange DE, van de Wetering M, and Clevers H. OLFM4 is a robust marker for stem cells in human intestine and marks a subset of colorectal cancer cells. Gastroenterology 137: 15–17, 2009.

68. Wang M, Liang C, Hu H, Zhou L, Xu B, Wang X, Han Y, Nie Y, Jia S, Liang J, and Wu K. Intraperitoneal injection (IP), Intravenous injection (IV) or anal injection (AI)? Best way for mesenchymal stem cells transplantation for colitis. Sci Rep 6: 30696, 2016.

69. Weil BR, Markel TA, Herrmann JL, Abarbanell AM, and Meldrum DR. Mesenchymal stem cells enhance the viability and proliferation of human fetal intestinal epithelial cells following hypoxic injury via paracrine mechanisms. Surgery 146: 190–197, 2009.

70. Weis VG, Petersen CP, Weis JA, Meyer AR, Choi E, Mills JC, and Goldenring JR. Maturity and age influence chief cell ability to transdifferentiate into metaplasia. Am J Physiol Gastrointest Liver Physiol 312: G67–G76, 2017.

71. Weis VG, Sousa JF, LaFleur BJ, Nam KT, Weis JA, Finke PE, Ameen NA, Fox JG, and Goldenring JR. Heterogeneity in mouse spasmolytic polypeptide-expressing metaplasia lineages identifies markers of metaplastic progression. Gut 62: 1270–1279, 2013.

72. Weitkamp JH, Koyama T, Rock MT, Correa H, Goettel JA, Matta P, Oswald-Richter K, Rosen MJ, Engelhardt BG, Moore DJ, and Polk DB. Necrotising enterocolitis is characterised by disrupted immune regulation and diminished mucosal regulatory (FOXP3)/effector (CD4, CD8) T cell ratios. Gut 62: 73–82, 2013.

73. Yang J, Watkins D, Chen CL, Bhushan B, Zhou Y, and Besner GE. Heparin-binding epidermal growth factor-like growth factor and mesenchymal stem cells act synergistically to prevent experimental necrotizing enterocolitis. J Am Coll Surg 215: 534–545, 2012.

74. Yang J, Watkins D, Chen CL, Zhang HY, Zhou Y, Velten M, and Besner GE. A technique for systemic mesenchymal stem cell transplantation in newborn rat pups. J Invest Surg 25: 405–414, 2012.

75. Yeh KY. Small intestine of artificially reared rat pups: effect of caloric intake and dietary composition on growth and disaccharidase activities. J Nutr 113: 1496–1502, 1983.

76. Yu R, Jiang S, Tao Y, Li P, Yin J, and Zhou Q. Inhibition of HMGB1 improves necrotizing enterocolitis by inhibiting NLRP3 via TLR4 and NF-kappaB signaling pathways. J Cell Physiol 234: 13431–13438, 2019.

77. Zani A, Cananzi M, Fascetti-Leon F, Lauriti G, Smith VV, Bollini S, Ghionzoli M, D’Arrigo A, Pozzobon M, Piccoli M, Hicks A, Wells J, Siow B, Sebire NJ, Bishop C, Leon A, Atala A, Lythgoe MF, Pierro A, Eaton S, and De Coppi P. Amniotic fluid stem cells improve survival and enhance repair of damaged intestine in necrotising enterocolitis via a COX-2 dependent mechanism. Gut 63: 300–309, 2014.

78. Zani A, Cananzi M, Lauriti G, Fascetti-Leon F, Wells J, Siow B, Lythgoe MF, Pierro A, Eaton S, and De Coppi P. Amniotic fluid stem cells prevent development of ascites in a neonatal rat model of necrotizing enterocolitis. Eur J Pediatr Surg 24: 57–60, 2014.

79. Zani A, Cordischi L, Cananzi M, De Coppi P, Smith VV, Eaton S, and Pierro A. Assessment of a neonatal rat model of necrotizing enterocolitis. Eur J Pediatr Surg 18: 423–426, 2008.

80. Zhang C, Sherman MP, Prince LS, Bader D, Weitkamp JH, Slaughter JC, and McElroy SJ. Paneth cell ablation in the presence of Klebsiella pneumoniae induces necrotizing enterocolitis (NEC)-like injury in the small intestine of immature mice. Dis Model Mech 5: 522–532, 2012.

81. Zhang H, Deng T, Liu R, Bai M, Zhou L, Wang X, Li S, Wang X, Yang H, Li J, Ning T, Huang D, Li H, Zhang L, Ying G, and Ba Y. Exosome-delivered EGFR regulates liver microenvironment to promote gastric cancer liver metastasis. Nat Commun 8: 15016, 2017.

